# Molecular details and phospho-regulation of the CENP-T-Mis12 complex interaction during mitosis in DT40 cells

**DOI:** 10.1101/2024.06.20.599825

**Authors:** Yusuke Takenoshita, Masatoshi Hara, Reiko Nakagawa, Mariko Ariyoshi, Tatsuo Fukagawa

## Abstract

To establish bi-polar attachments of microtubules to sister chromatids, an inner kinetochore subcomplex, the constitutive centromere-associated network (CCAN), is assembled on centromeric chromatin and recruits the microtubule-binding subcomplex called the KMN network. Although it is important to characterize the interaction between CCAN and KMN, it is difficult to evaluate the significance of each interaction in cells, since CCAN proteins CENP-C and CENP-T independently bind to the Mis12 complex (Mis12C) of KMN. Here, we demonstrate molecular details of the CENP-T-Mis12C interaction using chicken DT40 cells lacking CENP-C-Mis12C interaction. Based on AlphaFold2 predictions, we identified two binding surfaces for the CENP-T-Mis12C interaction, demonstrating that each surface is critical for Mis12C recruitment and cell viability. This interaction via two interaction surfaces is cooperatively regulated by dual phosphorylation of Dsn1 (a Mis12C component) and CENP-T, ensuring robust CENP-T-Mis12C interaction and proper mitotic progression. These findings deepen our understanding of kinetochores assembly in cells.

## Introduction

Chromosome segregation during mitosis is critical for transmitting genetic information to the progeny. This is achieved by attaching sister chromatids to bipolar mitotic spindles, leading to chromosome segregation into daughter cells. Spindle microtubules bind to a large protein complex called the kinetochore, which is formed on the centromere of each sister chromatid to ensure accurate chromosome segregation ^1–4^. To understand the mechanisms underlying chromosome segregation, it is crucial to clarify how the kinetochore is assembled on the centromere.

Kinetochores comprise two major complexes. One is the constitutive centromere-associated network (CCAN), containing 16 protein components that constitutively localize to the centromere throughout the cell cycle, forming a base for kinetochore assembly ^2,3,5–10^. Another large complex is the KMN network, comprising the Knl1 complex (Knl1C), Mis12 complex (Mis12C), and Ndc80 complex (Ndc80C), which are associated with CCAN from G_2_ phase to mitosis to establish a functional kinetochore structure for spindle microtubule binding ^2,6–8^. Ndc80C in the KMN network directly associates with spindle microtubules ^11–13^, and also associates with CCAN directly or via Mis12C ^2,7,8^. Since CCAN associates with centromeric chromatin, the linkage between CCAN and the KMN network is critical for bridging centromeric chromatin with spindle microtubules, ultimately ensuring accurate chromosome segregation.

In vertebrate cells, the KMN network is recruited to the CCAN through two parallel pathways: the CENP-C and CENP-T pathways ^1,3,4,6,14–19^. Mis12C binds to CENP-C and CENP-T independently in each pathway, acting as a hub for KMN network assembly by recruiting Ndc80C and Knl1C ^20^. In addition, CENP-T directly recruits Ndc80C via the Ndc80C binding domain(s) at its N-terminus (human CENP-T has two binding domains and chicken CENP-T has one domain) ^15,17,21^. Although both pathways contribute to Mis12C recruitment for KMN network assembly at an appropriate time ^16–18,22,23^, the CENP-T-Mis12C interaction has a dominant function in chromosome segregation compared to the CENP-C-Mis12C interaction ^18,23^. Previous studies have proposed a molecular basis for the CENP-T-Mis12C interaction based on *in vitro* binding assays and structural prediction analyses ^21,24,25^. However, it remains unclear how the CENP-T-Mis12C interaction is regulated and how this regulatory mechanism is significant in cells, as it is challenging to distinguish CENP-C-and CENP-T-binding Mis12C in native kinetochores. Therefore, clarifying the regulatory mechanisms and biological significance of CENP-T-Mis12C interaction in cells is not trivial.

In this study, we examined the molecular details of the CENP-T-Mis12C interaction at the native kinetochore using chicken DT40 mutant cells lacking the Mis12C binding region of CENP-C. Combined with structure prediction and cell biology analyses, we identified two binding sites in CENP-T for Mis12C interaction, which are essential for cell viability. We also propose that the dual phosphorylation of Dsn1 and CENP-T cooperatively contributes to the CENP-T-Mis12C interaction, ensuring mitotic progression in cells. The molecular details of the dual phospho-regulation of the CENP-T-Mis12C interaction provided insights into the robust assembly mechanism of kinetochores for accurate chromosome segregation.

## Results

### Localization profile of CENP-C-and CENP-T-binding Mis12C

Both CENP-C and CENP-T contain a binding region for Mis12C ^14,15,18,21,23,24,26,27^; however, it is unclear when and how Mis12C is recruited to each binding region. To distinguish the cell cycle timing of Mis12C recruitment to CENP-C and CENP-T, we examined Mis12C localization at kinetochores in CENP-C^WT^ cells (wild-type chicken DT40 cells) and CENP-C^Δ73^ cells lacking the Mis12C-binding region of CENP-C (aa 1–73; Figure 1A, S1A, and S1C). To visualize Mis12C, we introduced a GFP-Dsn1 construct (a Mis12C component) into the endogenous Dsn1 locus (Figure S1B and S1C). Consistent with our previous findings showing that Mis12C starts to localize to centromeres in interphase and continues to localize during mitosis in chicken DT40 cells ^18^, GFP-Dsn1 localized to centromeres in both interphase and M phase in CENP-C^WT^ cells (Figure 1B). However, GFP-Dsn1 localized to the centromeres only during mitosis and not during interphase in CENP-C^Δ73^ cells (Figure 1B), suggesting that Mis12C associates only with CENP-C and does not bind to CENP-T in interphase cells. Previous studies have suggested that the CENP-C interaction surface of Mis12C is masked by the Dsn1 basic motif to inhibit the CENP-C-Mis12C interaction during interphase, and this mask is precluded by phosphorylation of the basic motif by the mitotic kinase Aurora B during mitosis ^18,25,27,28^. However, Mis12C was recruited to CENP-C in the interphase of DT40 cells (Figure 1B). As discussed in a previous study ^25^, Mis12C may interact with CENP-C via an additional interaction surface that is not masked by the Dsn1 basic motif (Figure S1D‒S1I; see Discussion). Our results suggest that the additional interaction surface is used for CENP-C-Mis12C interaction in interphase cells.

**Figure 1.**
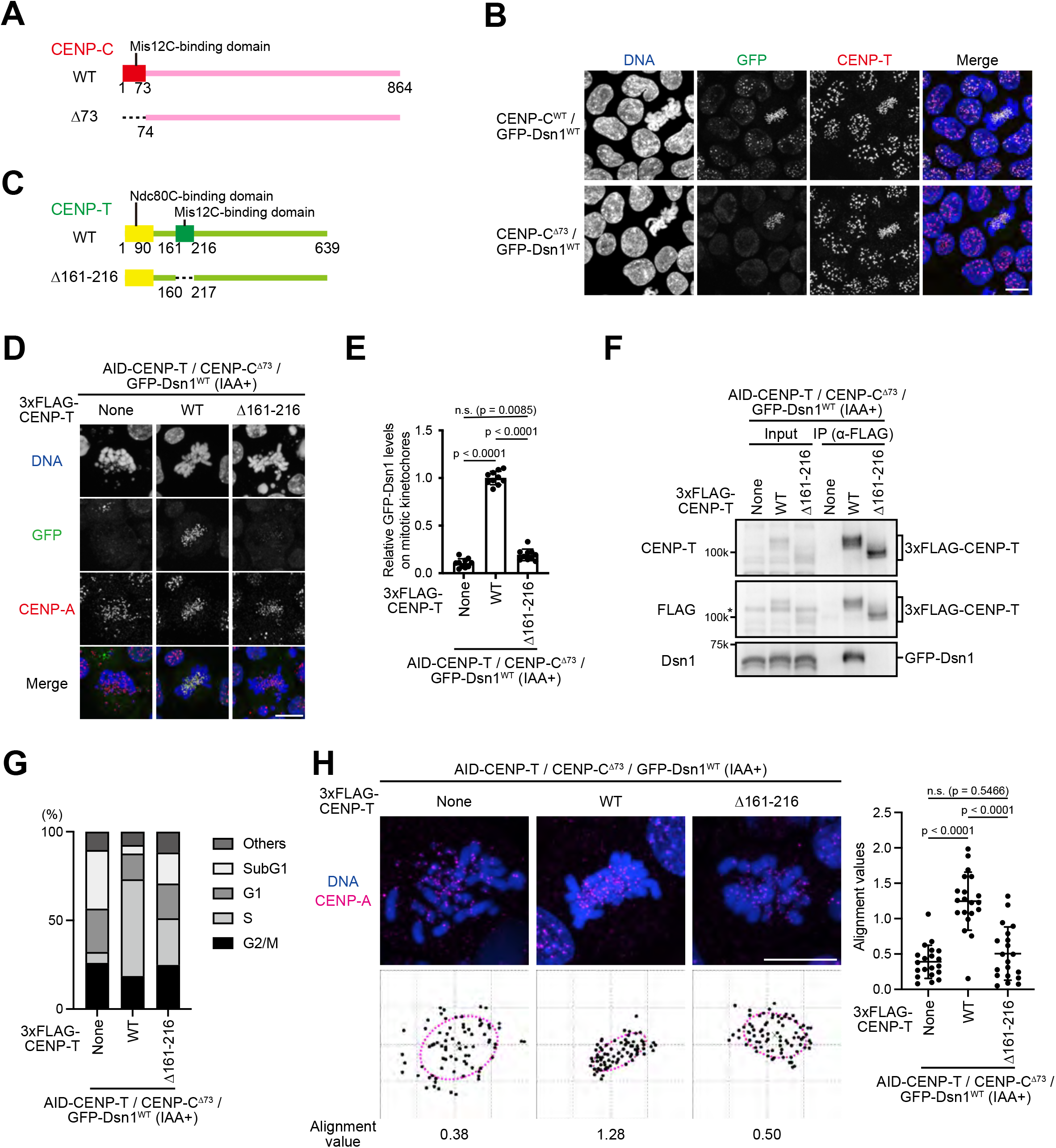
CENP-T interacts with Mis12C during mitosis. (A) Schematic representation of chicken CENP-C (864 aa). The Mis12C-binding domain was deleted in CENP-C^Δ73^. (B) Localization of GFP-Dsn1 in CENP-C^WT^ and CENP-C^Δ73^ cells. Cells were fixed and stained with an anti-CENP-T antibody. DNA was stained with DAPI. Scale bar, 10 µm. (C) Schematic representation of chicken CENP-T (639 aa). The Mis12C-binding domain was deleted in CENP-T^Δ161–216^. (D) Localization of GFP-Dsn1. 3xFLAG-CENP-T^WT^ or 3xFLAG-CENP-T^Δ161–216^ was expressed in AID-based CENP-T conditional knockdown CENP-C^Δ73^ cells in which wild-type CENP-T protein was degraded by IAA addition. Cells were treated with IAA for 12 h, fixed, and stained with an anti-CENP-A antibody. DNA was stained with DAPI. Scale bar, 10 µm. (E) Quantification data of GFP-Dsn1 signals at kinetochores in mitotic cells shown in (D). Error bars show the mean ± SD; *p*-values were calculated by one-way ANOVA (F (2,27) = 672.4; p<0.0001) followed by Tukey’s test. (F) Immunoprecipitation using anti-FLAG antibody in AID-based CENP-T conditional knockdown CENP-C^Δ73^/GFP-Dsn1 cells expressing 3xFLAG-CENP-T^WT^ or 3xFLAG-CENP-T^Δ161–216^. Cells were cultured with IAA for 12 h and nocodazole was added for last 10 h before immunoprecipitation experiments. Immunoprecipitated samples were subjected to immunoblot analyses using anti-CENP-T, anti-FLAG, and anti-Dsn1 antibodies. Asterisk indicates nonspecific bands. (G) Cell cycle distribution profiles of AID-based CENP-T conditional knockdown CENP-C^Δ73^/GFP-Dsn1 cells expressing 3xFLAG-CENP-T^WT^ or 3xFLAG-CENP-T^Δ161–216^ in the presence of IAA for 12 h. (H) Chromosome alignment values in AID-based CENP-T conditional knockdown CENP-C^Δ73^/GFP-Dsn1 cells expressing 3xFLAG-CENP-T^WT^ or 3xFLAG-CENP-T^Δ161–216^ in the presence of IAA for 12 h and MG132 for the last 2 h. Error bars show the mean ± SD; *p*-values were calculated by one-way ANOVA (F (2,57) = 35.49; p<0.0001) followed by Tukey’s test.

We previously showed that two CENP-T regions (aa 161–200 and aa 201–216) are required for interaction with Mis12C ^23^. To examine whether mitotic kinetochore localization of Mis12C in CENP-C^Δ73^ cells depends on the aa 161–216 region of CENP-T, we introduced a FLAG-fused CENP-T mutant construct lacking the aa 161–216 region into the endogenous CENP-T locus in CENP-C^Δ73^ cells expressing Auxin-Inducible Degron (AID)-tagged CENP-T (CENP-C^Δ73^/CENP-T^Δ161–216^ cells; Figure 1C and S2A and S2B). In this cell line, wild-type CENP-T was degraded after 3-indole acetic acid (IAA) addition, and instead of wild-type CENP-T, only CENP-T^Δ161–216^ was expressed (Figure S2C). As a control, the FLAG-fused wild-type CENP-T construct was introduced into the CENP-T locus (CENP-C^Δ73^/CENP-T^WT^ cells; Figure S2C). Next, we examined the mitotic localization of Mis12C in these cells. The mitotic GFP-Dsn1 localization observed in CENP-C^Δ73^/CENP-T^WT^ cells was almost undetectable in CENP-C^Δ73^/CENP-T^Δ161–216^ cells (Figure 1D and 1E). Consistent with this result, Mis12C co-immunoprecipitated with CENP-T^WT^ but not with CENP-T^Δ161–216^ (Figure 1F). In addition to the CENP-T-Mis12C interaction data, we examined the cell cycle profile of CENP-C^Δ73^/CENP-T^Δ161–216^ cells by flow cytometry. The G2/M and subG1 fractions were increased in CENP-C^Δ73^/CENP-T^Δ161–216^ cells, similar to AID-based CENP-T knockdown in CENP-C^Δ73^ cells (Figure 1G), suggesting that mitotic arrest and subsequent cell death occurred in CENP-C^Δ73^/CENP-T^Δ161–216^ cells. To further analyze the mitotic phenotype of CENP-C^Δ73^/CENP-T^Δ161–216^ cells, we examined the chromosome alignment of mitotic cells. As shown in Figure 1H, chromosomes were not aligned in CENP-C^Δ73^/CENP-T^Δ161–216^ cells and AID-based CENP-T knockdown CENP-C^Δ73^ cells, whereas chromosomes were well aligned in the metaphase plate in CENP-C ^Δ73^/CENP-T^WT^ cells. Taken together, we propose that the CENP-T aa 161– 216 region is required for Mis12C mitotic kinetochore localization via CENP-T-Mis12C interaction in CENP-C^Δ73^ cells and is critical for proper mitotic progression due to its binding to Mis12C.

### Structural prediction of interfaces for the CENP-T-Mis12C interaction

Next, to examine the molecular basis of the CENP-T-Mis12C interaction, we utilized AlphaFold2 to predict the structure of Mis12C, including four full-length proteins (Mis12, Dsn1, Nsl1, and Pmf1) in complex with the aa 161‒216 region of CENP-T (Figure 2A, Figure S3A‒S3D). The predicted structure of chicken Mis12C comprised two head domains (Head1 and Head2) connected to a coiled-coil stalk domain, similar to the crystal structures of human Mis12C and yeast Mis12C (Figure 2A) ^27,29^. The model of CENP-T^161–216^ contains two *α*-helices connected via an extended linker region.

**Figure 2.**
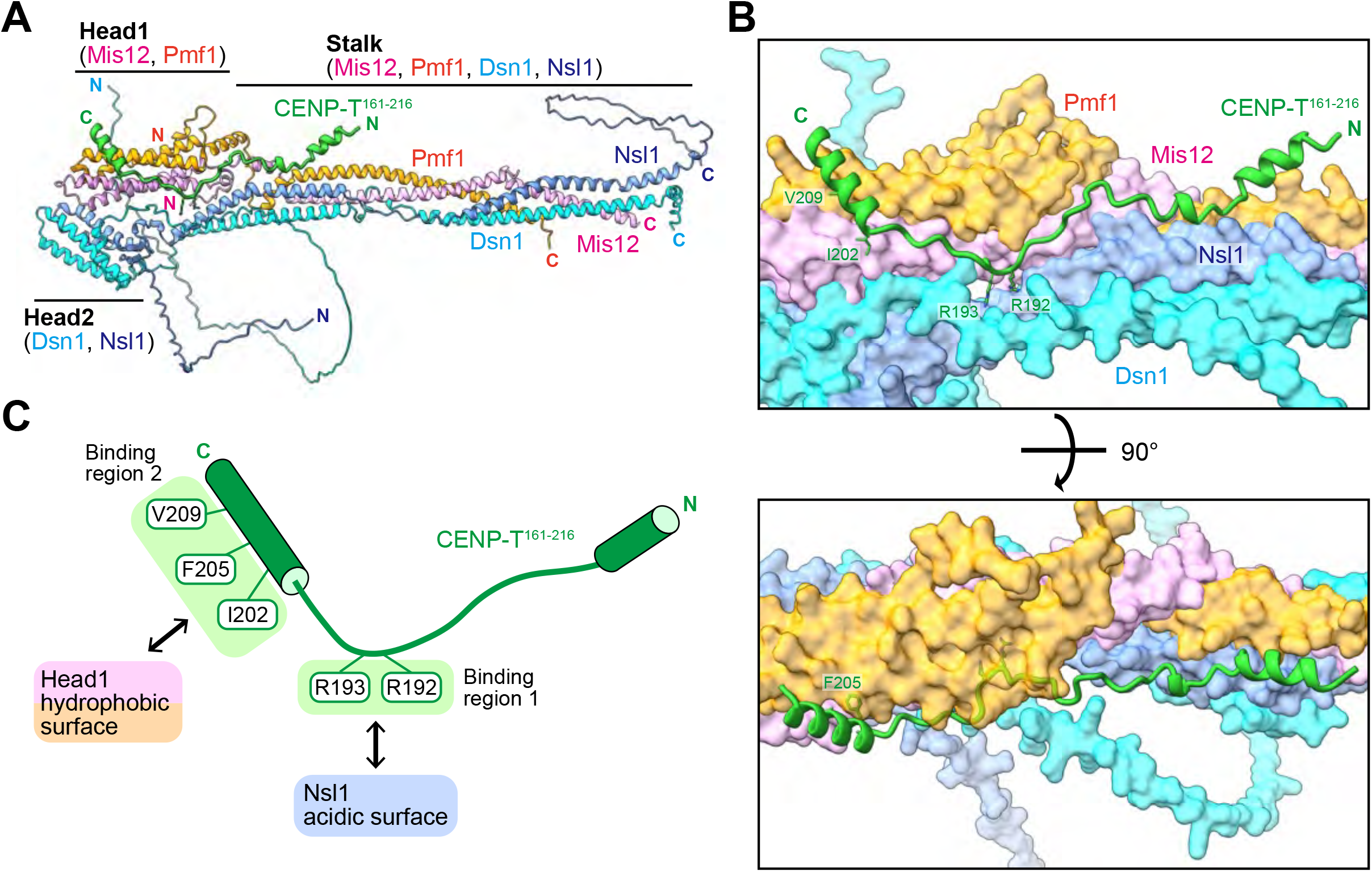
Molecular basis for the CENP-T-Mis12C interaction based on a complex structure predicted by AlphaFold2. (A) AlphaFold2 predicted structure of chicken Mis12C in complex with the Mis12C-binding domain of CENP-T (rank_1 model; rank_2, 3, 4, and 5 models are shown in Figure S3). The CENP-T aa 161–216 region and the full lengths of Mis12C components (Mis12, Pmf1, Dsn1, and Nsl1) were used for the prediction, displayed as a cartoon, and color-coded as indicated in the figure. (B) Overview of the CENP-T-Mis12C interface. Mis12C is shown as a surface model, and the CENP-T aa 161–216 region is shown as a cartoon. The side chains of key residues of CENP-T for binding to Mis12C are shown with labels. (C) Schematic representation of the CENP-T aa 161–216 structure. There are two binding regions for Mis12C (binding regions 1 and 2). Binding region 1 interacts with the acidic surface of Nsl1, and binding region 2 binds to the Head1 region of Mis12C via hydrophobic interactions.

AlphaFold2 predicted that CENP-T^161–216^ binds across the stalk domain to the Head1 domain of Mis12C (Figure 2A and 2B, Figure S3C and S3D). The predicted structure suggested two Mis12C-binding regions of CENP-T (binding region 1 and 2; Figure 2B and C). In the binding region 1, R192 and R193 of CENP-T in the extended region is supposed to interact with the Nsl1 acidic surface in the stalk region of Mis12C (Figure 2B and 2C). In the binding region 2, the C-terminal *α*-helix of CENP-T^161–216^ harboring I202, F206, and V209 is predicted to associate with the hydrophobic surface of the Head1 domain of Mis12C (Figure 2B and 2C). As these two Mis12C-binding regions of CENP-T seem important for the CENP-T-Mis12C interaction, we further analyzed the molecular details of the CENP-T-Mis12C interaction and its significance in CENP-C^Δ73^ cells.

### Electrostatic interaction of Mis12C with the binding region 1 of CENP-T

As described above, the AlphaFold2 model suggested that the two arginine residues of CENP-T (R192 and R193) in the binding region 1 form electrostatic interactions with the acidic surface of Nsl1 in Mis12C (Figure 3A). These arginine residues of CENP-T and the acidic region of Nsl1 are highly conserved (Figure S4A and S4B). To evaluate the importance of this interaction surface in chicken DT40 cells, we replaced R192 and R193 with alanine in CENP-T (CENP-T^RR-AA^) and introduced a mutant construct into the CENP-T locus in CENP-C^Δ73^ cells expressing AID-CENP-T (CENP-C^Δ73^/CENP-T^RR-AA^ cells; Figure S4C). We examined the GFP-Dsn1 levels at the mitotic kinetochores in these cells and found that these levels were ∼35% of those in CENP-C^Δ73^/CENP-T^WT^ cells, which were slightly higher than that in CENP-C^Δ73^/CENP-T^Δ161–216^ cells (Figure 3B and C). We also confirmed that Mis12C did not co-precipitate with CENP-T^RR-AA^ like CENP-T^Δ161–216^ (Figure 3D). Consistent with these results, we observed a clear growth delay in CENP-C^Δ73^/CENP-T^RR-AA^ cells following IAA addition (Figure S4D). Based on these results, we propose that CENP-T R192 and R193 form electrostatic interactions with the Nsl1 acidic region, essential for the CENP-T-Mis12C binding and cell proliferation.

**Figure 3.**
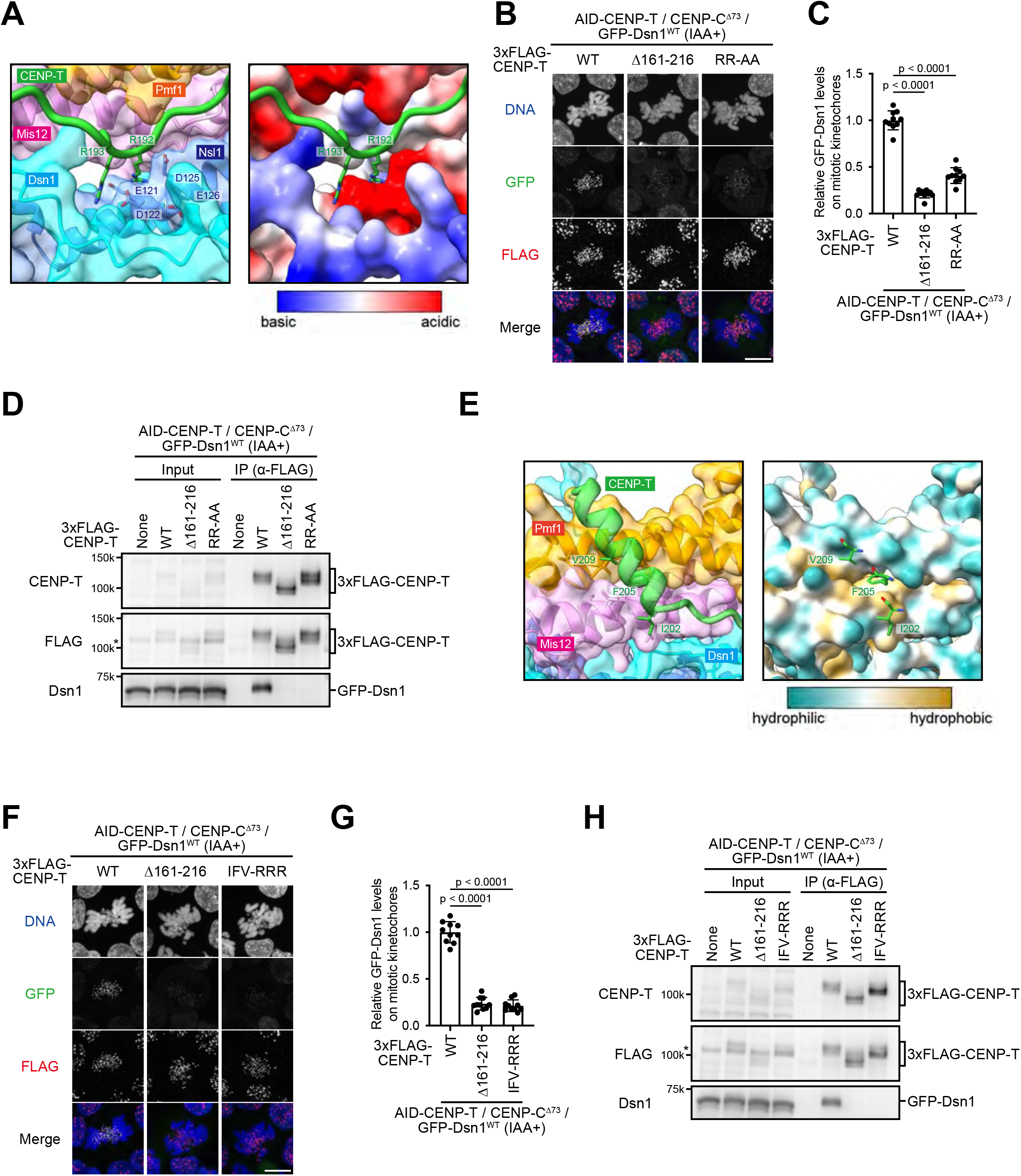
Two binding regions of CENP-T are required for the CENP-T-Mis12C interaction. (A) The magnified view of the Nsl1 acidic surface associated with the binding region 1 of CENP-T. Left: The transparent surface model of Mis12C is overlaid on a cartoon model. CENP-T is shown as a cartoon and models are color-coded as indicated in the figure. CENP-T R192 and R193 and Nsl1 acidic residues (E121, D122, D125, and E126) are labeled. Right: Electrostatic potential of Mis12C mapped on the surface representation. CENP-T is shown as a cartoon. (B) Localization of GFP-Dsn1. 3xFLAG-CENP-T^WT^, 3xFLAG-CENP-T^Δ161–216^, or 3xFLAG-CENP-T^RR-AA^ was expressed in AID-based CENP-T conditional knockdown CENP-C^Δ73^ cells in which wild-type CENP-T protein was degraded by IAA addition. Cells were treated with IAA for 12 h, fixed, and stained with an anti-FLAG antibody. DNA was stained with DAPI. Scale bar, 10 µm. (C) Quantification data of GFP-Dsn1 signals at kinetochores in mitotic cells shown in (B). Error bars show the mean ± SD; *p*-values were calculated by one-way ANOVA (F (2,27) = 262.6; p<0.0001) followed by Tukey’s test. (D) Immunoprecipitation using anti-FLAG antibody in AID-based CENP-T conditional knockdown CENP-C^Δ73^/GFP-Dsn1 cells expressing 3xFLAG-CENP-T^WT^, 3xFLAG-CENP-T^Δ161–216^, or 3xFLAG-CENP-T^RR-AA^. Cells were cultured with IAA for 12 h and nocodazole was added for last 10 h before immunoprecipitation experiments. Immunoprecipitated samples were subjected to immunoblot analyses using anti-CENP-T, anti-FLAG, and anti-Dsn1 antibodies. Asterisk indicates nonspecific bands. (E) Magnified view of the Mis12C-Head1 hydrophobic surface interacting with the binding region 2 of CENP-T. Left: The transparent surface model of Mis12C is overlaid on a cartoon model. CENP-T is shown as a cartoon and models are color-coded as indicated in the figure. CENP-T I202, F205, and V209 are labeled. Right: Electrostatic potential of Mis12C mapped on the surface representation. CENP-T is shown as a cartoon. (F) Localization of GFP-Dsn1. 3xFLAG-CENP-T^WT^, 3xFLAG-CENP-T^Δ161–216^, or 3xFLAG-CENP-T^IFV-RRR^ was expressed in AID-based CENP-T conditional knockdown CENP-C^Δ73^ cells. Cells were treated with IAA for 12 h, fixed, and stained with an anti-FLAG antibody. DNA was stained with DAPI. Scale bar, 10 µm. (G) Quantification data of GFP-Dsn1 signals at kinetochores in mitotic cells shown in (F). Error bars show the mean ± SD; *p*-values were calculated by one-way ANOVA (F (2,27) = 284.8; p<0.0001) followed by Tukey’s test. (H) Immunoprecipitation using anti-FLAG antibody in AID-based CENP-T conditional knockdown CENP-C^Δ73^/GFP-Dsn1 cells expressing 3xFLAG-CENP-T^WT^, 3xFLAG-CENP-T^Δ161–216^, or 3xFLAG-CENP-T^IFV-RRR^. Cells were cultured with IAA for 12 h and nocodazole was added for last 10 h before immunoprecipitation experiments. Immunoprecipitated samples were subjected to immunoblot analyses using anti-CENP-T, anti-FLAG, and anti-Dsn1 antibodies. Asterisk indicates nonspecific bands.

### Hydrophobic interaction of Mis12C with the binding region 2 of CENP-T

Structure prediction also suggested that the hydrophobic residues of CENP-T (I202, F205, and V209) aligning on the *α*-helix in the binding region 2 were associated with the hydrophobic surface of the Mis12C Head1 domain (Figure 3E). Although the hydrophobic residues of CENP-T are not thoroughly conserved in other species, the corresponding regions in human and mouse CENP-T contain a cluster of hydrophobic residues (Figure S4A). To evaluate the importance of this hydrophobic interface, we replaced I202, F205, and V209 of CENP-T with arginine (CENP-T^IFV-RRR^) and introduced a mutant construct into the CENP-T locus of CENP-C^Δ73^ cells expressing AID-fused CENP-T (CENP-C^Δ73^/CENP-T^IFV-RRR^ cells; Figure S4E). We observed a significant reduction in GFP-Dsn1 levels in the mitotic kinetochores of CENP-C^Δ73^/CENP-T^IFV-RRR^ cells compared to those of CENP-C^Δ73^/CENP-T^WT^ cells (Figure 3F and 3G), and Mis12C did not co-precipitate with CENP-T^IFV-RRR^ (Figure 3H). As in the case of CENP-C^Δ73^/CENP-T ^Δ161–216^ cells, after IAA addition, CENP-C^Δ73^/CENP-T^IFV-RRR^ cells stopped growing and ultimately died (Figure S4D). These results indicate that the hydrophobic interaction of the binding region 2 of CENP-T with Mis12C is essential for the CENP-T-Mis12C binding and cell viability. CENP-T has at least two binding regions for Mis12C, each necessary for binding to Mis12C. This result is consistent with our previous finding that both CENP-T aa 161–200 (containing binding region 1) and aa 201–216 (containing binding region 2) are required for CENP-T function via binding to Mis12C ^23^.

### Phosphorylation of Dsn1 in Mis12C facilitates stable CENP-T-Mis12C interaction

Although CENP-C interacts with Mis12C in both interphase and M phase, CENP-T interacts with Mis12C during mitosis but not in interphase (Figure 1). This implied that there are regulatory mechanisms by which CENP-T binds to Mis12C only during mitosis. The structural model of the human KMN network suggests that the Dsn1 basic motif binds to the Nsl1 acidic surface ^25,28^. The corresponding region of chicken Nsl1 is recognized by the CENP-T binding region 1 (Figure 3), and our AlphaFold2 prediction model for chicken Mis12C also suggests that the Dsn1 basic motif binds to the acidic surface of Nsl1 (Figure 4A). The Dsn1 basic motif possesses Aurora B phosphorylation sites ^16,30^, which inhibit the binding of the Dsn1 basic motif to the Nsl1 acidic surface ^25,27–29^. Thus, the interaction between the binding region 1 of CENP-T and the Nsl1 acidic surface might be inhibited by the Dsn1 basic motif during interphase; this inhibition is released during mitosis by Aurora B phosphorylation, similarly to the CENP-C-Mis12C interaction ^18,27,29^. Although this possibility was discussed based on an *in vitro* binding assay and structural model of human Mis12C ^24,25^, it is unclear whether this mechanism regulates the CENP-T-Mis12C interaction in cells and how it contributes to the function of the CENP-T pathway. To test this hypothesis, we constructed a Dsn1 construct lacking the basic motif (aa 93-114, Dsn1^Δ93–114^) and introduced it into the endogenous Dsn1 locus of CENP-C^Δ73^ cells (Figure 4B). While Dsn1^WT^ localized to kinetochores only during mitosis, Dsn1^Δ93–114^ localized to both interphase and mitotic kinetochores in CENP-C^Δ73^ cells (Figure 4C). Since both binding region 1 and 2 of CENP-T are essential for the Mis12C binding (Figure 3), CENP-T seems to interact with Mis12C via binding region 1 and 2 in CENP-C^Δ73^/Dsn1^Δ93–114^ interphase cells. Combined, we propose that the primary motif masks the binding surface of Mis12C for binding region 1 of CENP-T, and the mask is released upon phosphorylation, leading to the interaction between binding region 1 of CENP-T and Mis12C during mitosis. In turn, the release of the mask also facilitates the interaction between binding region 2 of CENP-T and Mis12C (Figure S5B). These results suggest a linkage between the regulatory mechanisms governing the interactions of binding region 1 and 2 of CENP-T with Mis12C.

**Figure 4.**
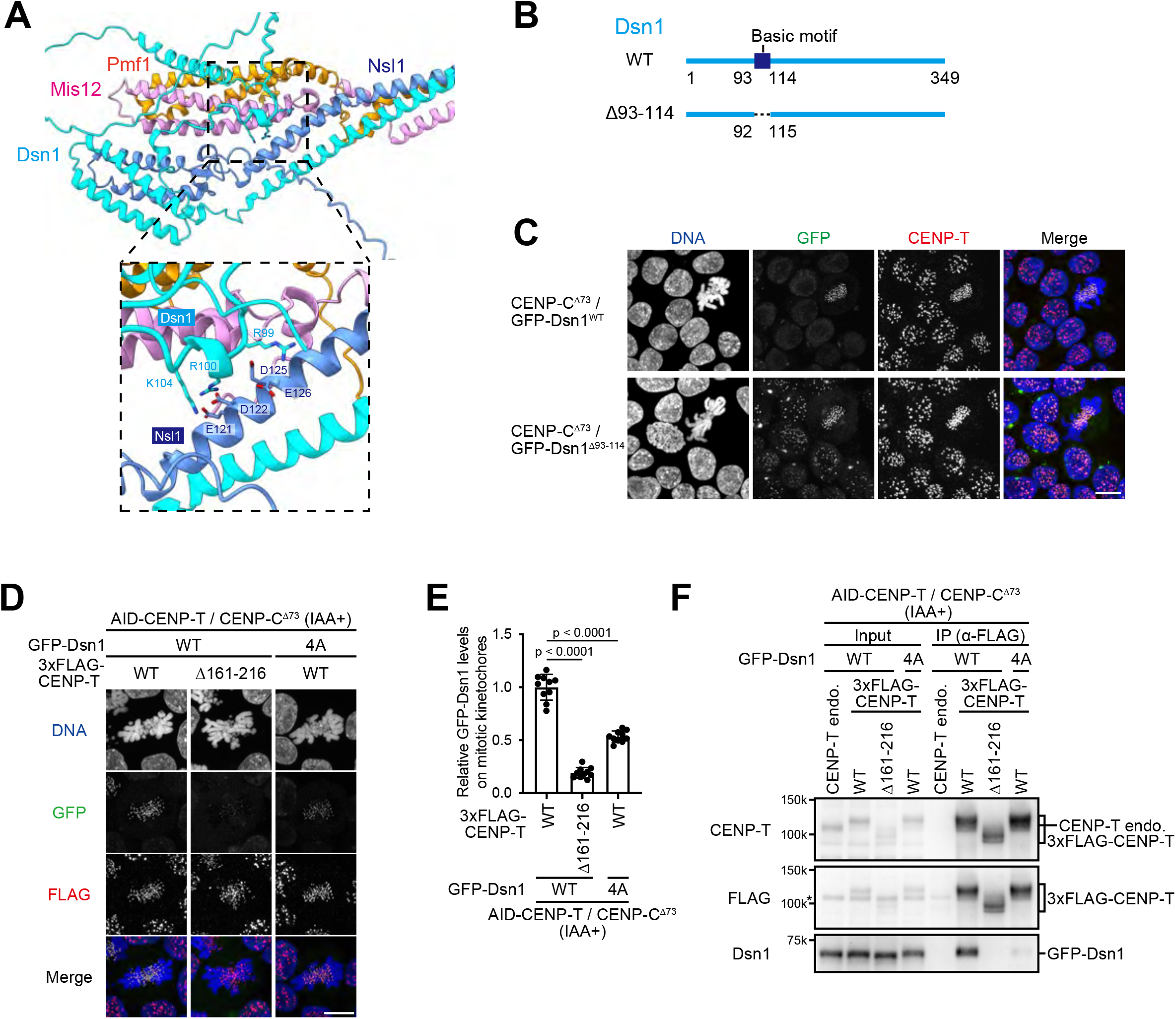
Dsn1 basic motif regulates the CENP-T-Mis12C interaction. (A) Overview of the Dsn1 basic motif, which masks the Nsl1 acidic surface. Mis12C is shown as a cartoon and color-coded, as indicated in the figure. The key residues of the Dsn1-Nsl1 interaction are labeled. (B) Schematic representation of chicken Dsn1 (349 aa). The basic motif was deleted in Dsn1^Δ93–114^. (C) Localization of GFP-Dsn1^WT^ or GFP-Dsn1^Δ93–114^ in CENP-C^Δ73^ cells. Cells were fixed and stained with an anti-CENP-T antibody. DNA was stained with DAPI. Scale bar, 10 µm. (D) Localization of GFP-Dsn1. GFP-Dsn1^WT^ or GFP-Dsn1^4A^ was expressed in AID-based CENP-T conditional knockdown CENP-C^Δ73^/CENP-T^WT^ cells or CENP-C^Δ73^/CENP-T^Δ161–216^ cells. Cells were treated with IAA for 12 h, fixed, and stained with an anti-FLAG antibody. DNA was stained with DAPI. Scale bar, 10 µm. (E) Quantification data of GFP-Dsn1 signals at kinetochores in mitotic cells shown in (D). Error bars show the mean ± SD; *p*-values were calculated by one-way ANOVA (F (2,27) = 245.1; p<0.0001) followed by Tukey’s test. (F) Immunoprecipitation using anti-FLAG antibody in AID-based CENP-T conditional knockdown CENP-C^Δ73^/CENP-T^WT^ cells or CENP-C^Δ73^/CENP-T^Δ161–216^ cells expressing GFP-Dsn1^WT^ or GFP-Dsn1^4A^. Cells were cultured with IAA for 12 h and nocodazole was added for last 10 h before immunoprecipitation experiments. Immunoprecipitated samples were subjected to immunoblot analyses using anti-CENP-T, anti-FLAG, and anti-Dsn1 antibodies. Asterisk indicates nonspecific bands.

Next, we generated Dsn1^4A^, in which the Aurora B phosphorylation sites of Dsn1 were substituted with alanine and introduced into the endogenous Dsn1 locus in CENP-C^Δ73^ cells expressing AID-fused CENP-T (CENP-C^Δ73^/Dsn1^4A^ cells; Figure S5A and S5C). We quantified the Dsn1 levels in these cells and found that they were reduced to approximately 55% relative to CENP-C^Δ73^/Dsn1^WT^ cells (Figure 4D and 4E). We also demonstrated that Dsn1^4A^ did not co-precipitate well with CENP-T compared with Dsn1^WT^ (Figure 4F). The growth of CENP-C^Δ73^/Dsn1^4A^ cells was comparable to that of CENP-C^Δ73^/Dsn1^WT^ cells (Figure S5D), suggesting that half of the Dsn1 was sufficient for the viability of CENP-C ^Δ73^ cells. As the expression of Dsn1^4A^ did not completely suppress Mis12C localization in CENP-C^Δ73^ cells unlike the expression of CENP-T^RR-AA^ (with mutations in binding region 1 of CENP-T), there should be an additional regulatory mechanism for the CENP-T-Mis12C interaction.

### CENP-T phosphorylation is involved in the regulation of the CENP-T-Mis12C interaction

As an additional regulatory mechanism for the CENP-T-Mis12C interaction, CENP-T-mediated phosphorylation is one of the candidates because chicken CENP-T is highly phosphorylated during mitosis ^23^. First, we examined the phosphorylation sites in the CENP-T aa 161–216 region using mitotic DT40 cell extracts by mass spectrometry and found that S175 and T184 were phosphorylated during mitosis (Figure 5A and S6A). T184 in chicken CENP-T might correspond to S201 of human CENP-T, which is phosphorylated by CDK1 ^21,24^; however, surrounding sequences of S175 in chicken CENP-T are not clearly conserved in other species (Figure S4A). As these two residues were not included in the binding regions 1 and 2, we re-evaluated the AlphaFold 2 prediction model of Mis12C in complex with the CENP-T aa 161–216 region. As shown in Figure 5B, S175 and T184 were close to the basic regions of Nsl1 and Pmf1, suggesting that phosphorylation of these CENP-T residues might provide the additional Mis12C binding site mediated via electrostatic interactions and stabilize the CENP-T-Mis12C interface. We generated a CENP-T mutant construct in which S175 and T184 were substituted with alanine and introduced into the CENP-T locus in CENP-C^Δ73^ cells expressing AID-fused CENP-T (CENP-C^Δ73^/CENP-T^2A^ cells; Figure S6B). We examined GFP-Dsn1 levels at kinetochores in CENP-C^Δ73^/CENP-T^2A^ cells and found that GFP-Dsn1 levels were significantly reduced compared to those in CENP-C^Δ73^/CENP-T^WT^ cells (∼70% of those in CENP-C^Δ73^/CENP-T^WT^ cells; Figure 5C and 5D). We also found that the levels of Mis12C that co-precipitated with CENP-T^2A^ were lower than those with CENP-T^WT^ (Figure 5E). Based on these results, we propose that CENP-T phosphorylation contributed to regulating the CENP-T-Mis12 interaction (Figure S6D; see discussion). However, as in the case of CENP-C^Δ73^/Dsn1^4A^ cells, substantial growth defects were not observed in CENP-C^Δ73^/CENP-T^2A^ cells (Figure S6C), indicating that CENP-C ^Δ73^ cells were still viable even when the interaction between CENP-T phospho-sites and Mis12C was impaired.

**Figure 5.**
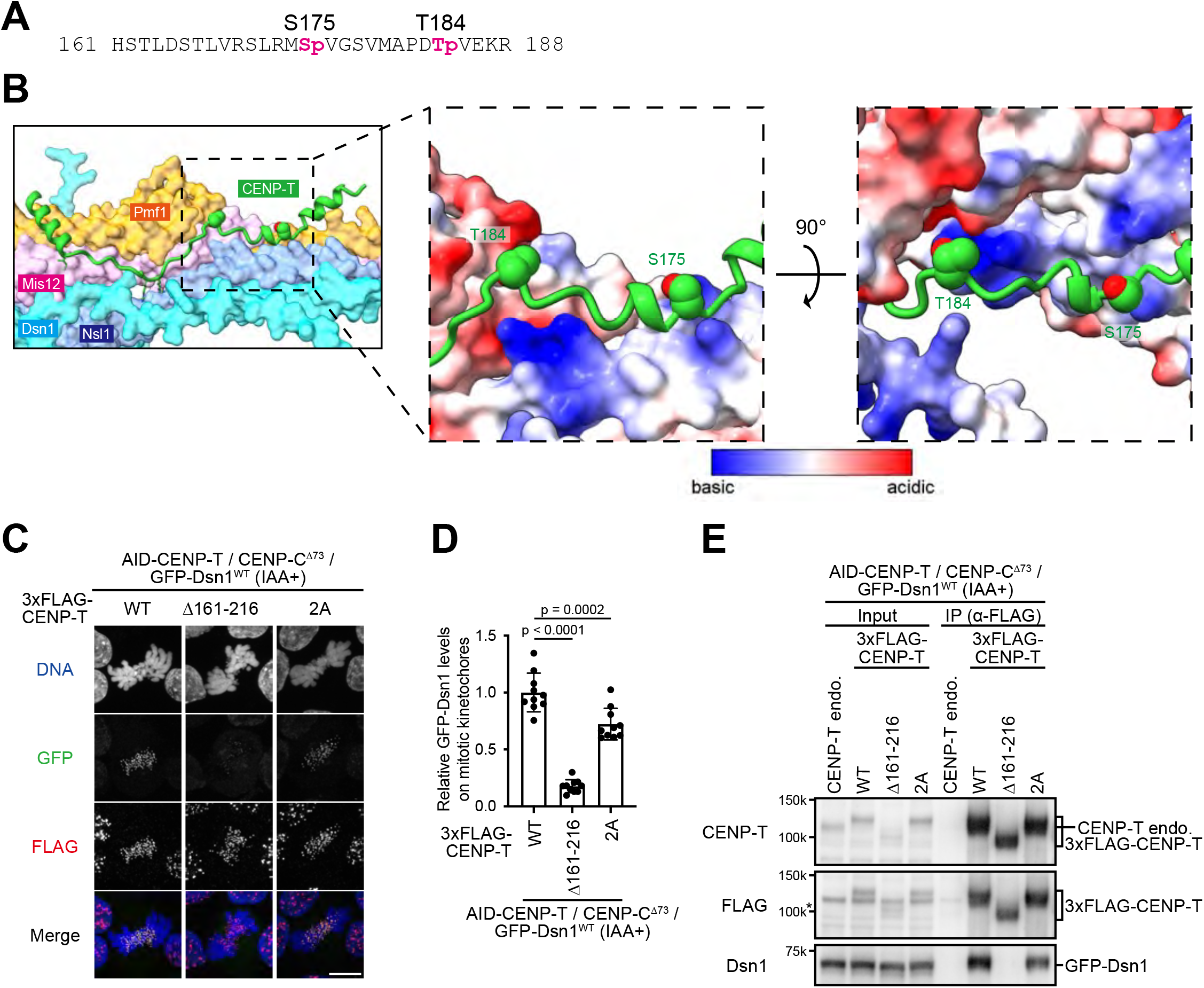
Phosphorylation of CENP-T regulates the CENP-T-Mis12C interaction. (A) S175 and T184 of CENP-T are phosphorylated in mitotic DT40 cells based on phospho-mass spectrometry analysis. (B) AlphaFold2 predicted the structure of chicken Mis12C in complex with the Mis12C-binding domain of CENP-T. Mis12C is shown as a surface model, and CENP-T is shown as a cartoon. S175 and T184 of CENP-T are labeled. The dashed box shows the electrostatic potential of Mis12C mapped on the surface representation. CENP-T is also shown as a cartoon. (C) Localization of GFP-Dsn1. 3xFLAG-CENP-T ^WT^, 3xFLAG-CENP-T^Δ161–216^, or 3xFLAG-CENP-T^2A^ was expressed in AID-based CENP-T conditional knockdown CENP-C^Δ73^ cells. Cells were treated with IAA for 12 h, fixed, and stained with an anti-FLAG antibody. DNA was stained with DAPI. Scale bar, 10 µm. (D) Quantification data of GFP-Dsn1 signals at kinetochores in mitotic cells shown in (C). Error bars show the mean ± SD; *p*-values were calculated by one-way ANOVA (F (2,27) = 102.4; p<0.0001) followed by Tukey’s test. (E) Immunoprecipitation using anti-FLAG antibody in AID-based CENP-T conditional knockdown GFP-Dsn1/CENP-C^Δ73^ cells expressing 3xFLAG-CENP-T^WT^, 3xFLAG-CENP-T^Δ161–216^, or 3xFLAG-CENP-T^2A^. Cells were cultured with IAA for 12 h and nocodazole was added for last 10 h before immunoprecipitation experiments. Immunoprecipitated samples were subjected to immunoblot analyses using anti-CENP-T, anti-FLAG, and anti-Dsn1 antibodies. Asterisk indicates nonspecific bands.

### Double mutant cells expressing CENP-T^2A^ and Dsn1^4A^ cause a more substantial reduction of Mis12C at kinetochores than single mutant cells

Although phosphorylation of Dsn1 and CENP-T facilitates the CENP-T-Mis12C interaction at different binding interfaces, inhibition of either phosphorylation did not impair CENP-C^Δ73^ cell proliferation. This prompted us to examine the phenotype of double mutant cells expressing CENP-T^2A^ and Dsn1^4A^. We introduced a construct containing either GFP-Dsn1^WT^ or GFP-Dsn1^4A^ into the endogenous Dsn1 locus of CENP-C^Δ73^/CENP-T^2A^ cells (Figure S7A). We then analyzed the effect of CENP-T and Dsn1 double mutations on Mis12C localization at the kinetochores. GFP-Dsn1 levels at kinetochores in CENP-C^Δ73^/CENP-T^2A^/Dsn1^WT^ and CENP-C^Δ73^/CENP-T^WT^/Dsn1^4A^ cells were reduced by ∼70% and ∼60%, respectively, compared to those in CENP-C^Δ73^/CENP-T^WT^/Dsn1^WT^ cells (Figure 6A and 6B). However, Dsn1 levels at kinetochores in CENP-C^Δ73^/CENP-T^2A^/Dsn1^4A^ cells were reduced to approximately 25% of those in CENP-C^Δ73^/CENP-T^WT^/Dsn1^WT^ cells (Figure 6A and 6B), indicating an additive reduction effect of Dsn1^4A^ and CENP-T^2A^ on Mis12C levels at kinetochores. This additive effect was confirmed using co-immunoprecipitation (Figure 6C). GFP-Dsn1 levels were further decreased in the CENP-T-immunoprecipitated fraction of CENP-C^Δ73^/CENP-T^2A^/Dsn1^4A^ cells, compared to those in the CENP-C^Δ73^/CENP-T^WT^/Dsn1^4A^ or CENP-C^Δ73^/CENP-T^2A^/Dsn1^WT^ cells (Figure 6C). Consistent with these results, CENP-C^Δ73^/CENP-T^2A^/Dsn1^4A^ cells showed a growth delay (Figure S7B). Then, we examined the M phase proportion in CENP-C^Δ73^/CENP-T^2A^/Dsn1^4A^ cells and found that the M phase proportion in CENP-C^Δ73^/CENP-T^2A^/Dsn1^4A^ cells was significantly increased compared to that in CENP-C^Δ73^/CENP-T^WT^/Dsn1^WT^, CENP-C^Δ73^/CENP-T^WT^/Dsn1^4A^, or CENP-C^Δ73^/CENP-T^2A^/Dsn1^WT^ cells (Figure 6D). We also assessed the sensitivity of these cells to low doses of nocodazole, inhibitor of microtubule polymerization. While the 5 ng/ml nocodazole treatment did not affect the cell viability of CENP-C^Δ73^/CENP-T^WT^/Dsn1^WT^, CENP-C^Δ73^/CENP-T^WT^/Dsn1^4A^, or CENP-C^Δ73^/CENP-T^2A^/Dsn1^WT^ cells, the cell viability of CENP-C^Δ73^/CENP-T^2A^/Dsn1^4A^ cells was decreased by the treatment with the same concentration of nocodazole (Figure 6E and F), suggesting that CENP-C^Δ73^/CENP-T^2A^/Dsn1^4A^ cells are sensitive to the low-dose nocodazole treatment possibly due to the reduction in the kinetochore-localized Mis12C levels. Based on these results, phosphorylation of both Dsn1 and CENP-T is important for timely and stable CENP-T-Mis12C interaction during mitosis, and this regulatory mechanism is critical for mitotic progression and CENP-C^Δ73^ cell proliferation.

**Figure 6.**
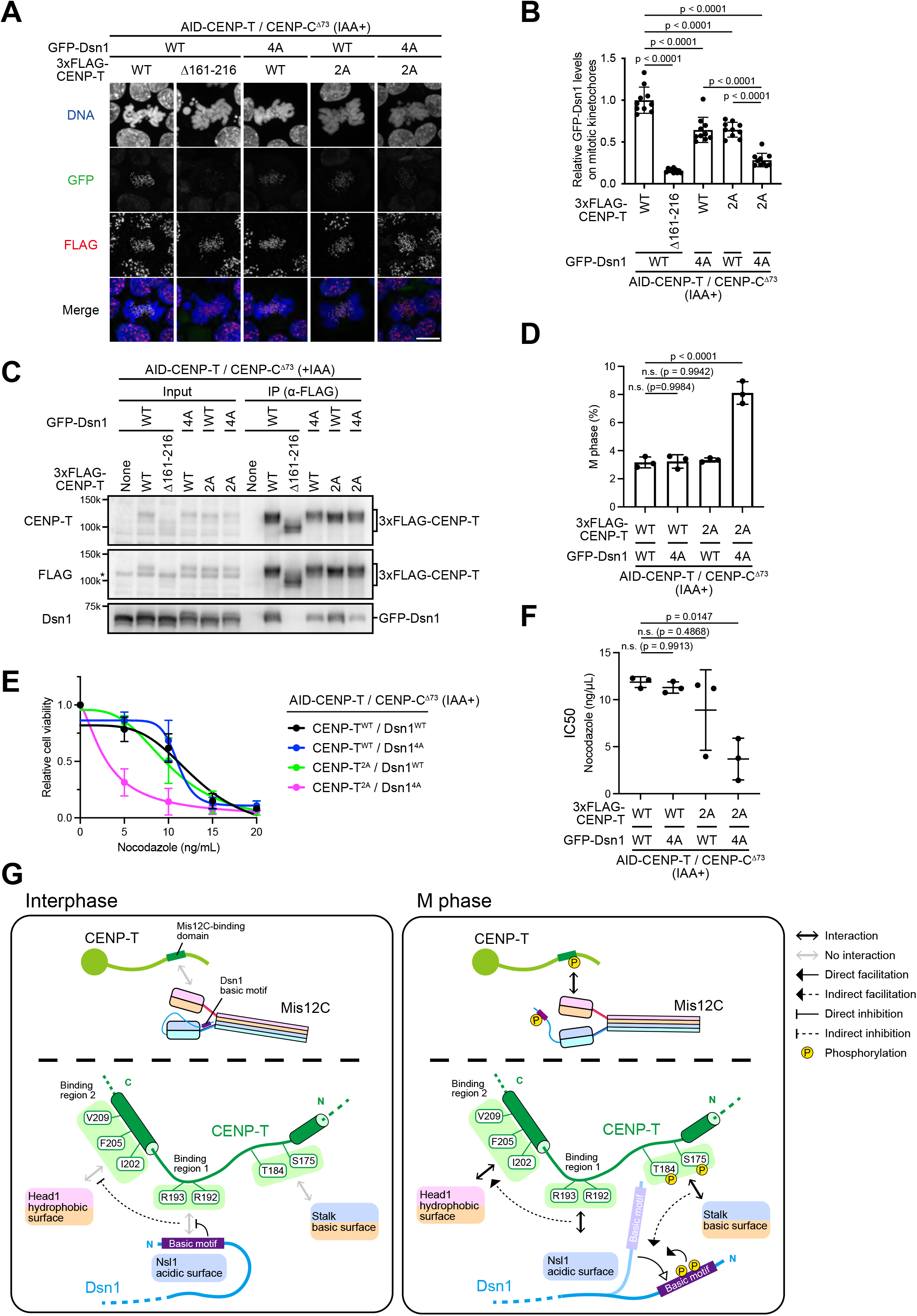
Dual phosphorylation of Dsn1 and CENP-T ensures the timely and robust CENP-T-Mis12C interaction. (A) Localization of GFP-Dsn1: 3xFLAG-CENP-T ^WT^, 3xFLAG-CENP-T^Δ161–216^, or 3xFLAG-CENP-T^2A^ was expressed in AID-based CENP-T conditional knockdown CENP-C^Δ73^/Dsn1^WT^ cells or CENP-C^Δ73^/Dsn1^4A^ cells. Cells were treated with IAA for 12 h, fixed, and stained with an anti-FLAG antibody. DNA was stained with DAPI. Scale bar, 10 µm. (B) Quantification of GFP-Dsn1 signals at kinetochores in mitotic cells shown in (A) Error bars show the mean ± SD; *p*-values were calculated by one-way ANOVA (F (4,45) = 89.33; p<0.0001) followed by Tukey’s test. (C) Immunoprecipitation using anti-FLAG antibody in AID-based CENP-T conditional knockdown CENP-C^Δ73^/ Dsn1^WT^ cells or CENP-C^Δ73^/Dsn1^4A^ cells expressing 3xFLAG-CENP-T^WT^, 3xFLAG-CENP-T^Δ161–216^, or 3xFLAG-CENP-T^2A^. Cells were cultured with IAA for 12 h and nocodazole was added for last 10 h before immunoprecipitation experiments. Immunoprecipitated samples were subjected to immunoblot analyses using anti-CENP-T, anti-FLAG, and anti-Dsn1 antibodies. Asterisk indicates nonspecific bands. (D) The ratio of M phase population in indicated cell lines. Cells were treated with IAA for 12 h and fixed. DNA was stained with DAPI. Populations for M phase and interphase were counted in each cell line, and the ratio of M phase cells was calculated. Error bars show the mean ± SD; *p*-values were calculated using one-way ANOVA (F (3,8) = 67.87; p<0.0001) followed by Tukey’s test. (E) Viability of cells treated with various nocodazole concentrations in indicated cell lines. 3xFLAG-CENP-T ^WT^ or 3xFLAG-CENP-T^2A^ was expressed in AID-based CENP-T conditional knockdown CENP-C^Δ73^/Dsn1^WT^ cells or CENP-C^Δ73^/Dsn1^4A^ cells, treated with IAA for 12 h, and added nocodazole. Cell viability was measured 24 h after nocodazole addition. Three independent experiments were performed. Error bars show the mean ± SD. (F) Half-maximal inhibitory concentration (IC50) of nocodazole calculated using the data shown in (E). Error bars show the mean ± SD; *p*-values were calculated by one-way ANOVA (F (3,8) = 6.965; p=0.0128) followed by Tukey’s test. (G) A model of the regulatory mechanism of the CENP-T-Mis12C interaction. In interphase, the Dsn1 basic motif masks the Nsl1 acidic surface, inhibiting the interaction between binding region 1 of CENP-T and the Nsl1 acidic surface of Mis12C. This inhibition leads to the inhibition of the interaction between binding region 2 of CENP-T and the Head1 hydrophobic surface of Mis12C. Since CENP-T is not phosphorylated in interphase, S175 and T184 of CENP-T do not interact with the stalk basic surface of Mis12C. In M phase, the Dsn1 basic motif is precluded from the Nsl1 acidic surface by the CENP-T and Dsn1 dual phospho-regulation. Phosphorylation of the Dsn1 basic motif by Aurora B preclude the mask by inhibiting the basic motif binding to the Nsl1 acidic surface. Phosphorylation of CENP-T facilitates to preclude the mask by unknown mechanisms. By coordination of CENP-T and Dsn1 dual phosphorylation, the Nsl1 acidic surface associates with binding region 1 of CENP-T. Additionally, CENP-T phosphorylation sites may directly bind to the Mis12C stalk basic domain.

## Discussion

In this study, we demonstrated that CENP-T binds to Mis12C via two interaction regions (binding regions 1 and 2), and each of them is essential for Mis12C binding. These interactions are cooperatively regulated by dual phosphorylation of Dsn1 and CENP-T, ensuring timely and robust CENP-T-Mis12C interactions during mitosis (Figure 6G; see below).

The two Mis12C-binding regions of CENP-T are essential for the CENP-T-Mis12C interaction. This makes it possible that the inhibition of the interaction of the binding region 1 of CENP-T with Mis12C regulated by the Dsn1 basic motif causes the defect of the interaction between binding region 2 of CENP-T and Mis12C. This mechanism results in complete inhibition of the CENP-T-Mis12C interaction in interphase (Interphase, Figure 6G). In addition, Mis12C containing the Dsn1 mutant lacking the basic motif (Dsn1^Δ93–114^) binds to CENP-T in interphase cells, suggesting that the deletion of the Dsn1 basic motif facilitates the interactions via two interaction surfaces. Therefore, we propose that phosphorylation of the Dsn1 basic motif by Aurora B can facilitate the interactions via two interaction surfaces simultaneously for timely CENP-T-Mis12C interaction during mitosis (M phase, Figure 6G).

A highlight of this paper is CENP-T and Dsn1 dual phospho-regulation for the CENP-T-Mis12C interaction during mitosis. Although each phosphorylation can recruit sufficient levels of Mis12C for cell proliferation, both mutations for CENP-T and Dsn1 showed additional reduction of Mis12C and caused mitotic defects. How are these dual phospho-regulation related each other? Mis12C levels were reduced in CENP-C^Δ73^/Dsn1^4A^ cells, but these are still higher than those in CENP-C^Δ73^/CENP-T^RR-AA^ cells that have mutations in binding region 1 of CENP-T, which is essential for Mis12C-binding. This result suggests that the mask of Dsn1 basic motif is precluded in some Mis12C populations, even if Aurora B phosphorylation sites of Dsn1 were mutated. We propose that the CENP-T phosphorylation contributes the exclusion of the Dsn1 basic motif mask by unknown mechanisms (Figure 6G and S6D). Additionally, the CENP-T phosphorylation sites may directly bind to Mis12C in parallel with phosphorylation of the Dsn1 basic motif. In any case, our results suggest that phosphorylation of CENP-T and Dsn1 can coordinately preclude the Dsn1 basic motif mask. Thus, in our model, the CENP-T and Dsn1 phosphorylation may contribute to an initiation of the CENP-T-Mis12C interaction in early mitosis (Figure S7D). By this initiation step, CENP-T starts to associate with Mis12C. Then, the Dsn1 mask is released by co-ordination of Dsn1 and CENP-T phosphorylation, and CENP-T-binding region 1 can stably bind to Nsl1 of Mis12C and subsequently hydrophobic interaction via CENP-T-binding region 2 with Mis12C can be established (Figure 6G and Figure S7D).

Interestingly, Mis12C localizes to interphase centromeres depends on CENP-C (Figure 1). For the CENP-C-Mis12C interaction during mitosis, the involvement of phospho-regulation of Dsn1 has been well studied ^18,25,27–29^. Our AlphaFold2 prediction suggests that CENP-C is likely to bind to Pmf1 and Mis12 (Figure S1I) during interphase, independent on the Dsn1 phospho-regulation. This finding supports the results of a previous *in vitro* binding assay showing that CENP-C binds to Mis12C in an auto-inhibited state via the Dsn1 basic motif ^25^. However, the significance of interphase Mis12C localization at kinetochores via CENP-C remains unclear. A recent study demonstrated that Mis12C constitutively localizes to the centromeres in mouse germ cells expressing the Dsn1 isoform, which lacks the basic motif ^31^. They proposed that the continuous kinetochore localization of Mis12C might be important for stable KMN network assembly because germ cells immediately enter meiosis II after meiosis I without DNA replication. Similarly, interphase Mis12C localization at kinetochores via CENP-C may be important for timely KMN network assembly for the subsequent mitosis in cells.

In both CENP-C-and CENP-T-pathways, phosphorylation by Aurora B facilitates the CENP-C-or CENP-T-Mis12C interaction. However, Aurora B also phosphorylates Ndc80C on kinetochores on unaligned chromosomes to correct erroneous microtubule attachments and phosphorylation levels by Aurora B is decreased at kinetochores on aligned chromosomes ^32^. However, even at Aurora B less-kinetochores on aligned chromosomes, Mis12C localizes at kinetochores. Therefore, we propose that the CENP-C-or CENP-T-Mis12C interaction independent of Aurora B-mediated regulations is important to maintain Mis12C at kinetochores on aligned chromosomes for proper mitotic progression.

Recent structural analyses of the KMN network or CCAN using *in vitro* reconstitution have provided valuable insights into the mechanism underlying kinetochore assembly ^9,10,25,28^. However, it is still largely unknown how kinetochore components play a role in mitotic progression. Therefore, it is critical to evaluate the significance of the binding surfaces between kinetochore components, which have been clarified by structural analyses. However, some kinetochore components have similar functions and simultaneously recruit the same proteins ^14,15,17,33^. This makes it difficult to investigate the significance of each interaction in cells. In this study, using chicken DT40 cells lacking the CENP-C-Mis12C interaction, we clearly elucidated the significance of the interaction surfaces and phospho-regulation for the CENP-T-Mis12C interaction in cells. This knowledge provides important insights into how functional kinetochores are established in cells.

## Methods

### Cell culture

Chicken DT40 cells ^34^ were cultured in Dulbecco’s modified Eagle’s medium (Nacalai Tesque) containing 10% fetal bovine serum (Sigma), 1% chicken serum (Thermo Fisher Scientific), 10 µM 2-mercaptoethanol (Sigma), and penicillin (100 units/mL)-streptomycin (100 µg/mL) (Thermo Fisher Scientific) at 38.5°C with 5% CO_2_. For degradation of mScarlet-mAID-CENP-T in AID based-CENP-T conditional knockdown cells, 500 µM 3-indole acetic acid (IAA, Wako) was added.

### Plasmid constructions

GFP-Dsn1^WT^, GFP-Dsn1^4A^, and GFP-Dsn1^Δ93–114^ ^18^, were cloned into the plasmid containing approximately 2 kb homology arm around the start codon of *Dsn1* with the puromycin resistance gene cassette and EcoGPT gene cassette by In-Fusion^®^ HD (TAKARA) (pBS_Dsn1-N KI_GFP-Dsn1^WT^, _GFP-Dsn1^4A^, or _GFP-Dsn1^Δ93–114^_PuroR or _EcoGPT).

cDNA for CENP-T full-length sequence was cloned into pAID1.2-CMV-NmScarlet-mAID (Addgene; #140618). To express the OsTIR1-T2A-BSR and mScarlet-mAID-CENP-T under control of the *CMV* promoter from the *PGK* locus, CMV promoter_OsTIR1-T2A-BSR_IRES_mScarlet-mAIDCENP-T fragment was cloned into the plasmid containing approximately 2 kb genome homology arm around the start codon of *PGK* by In-Fusion^®^ HD (TAKARA) (pBS_PGK N-KI_ pCMV-OsTIR1-T2A-BSR-IRES-mScarlet-mAID-CENP-T).

cDNA for CENP-T full-length sequence was cloned into p3xFLAG-CMV-10 (SIGMA). Several CENP-T mutants (CENP-T^Δ161–216^, CENP-T^RR-AA^, CENP-T^IFV-RRR^, and CENP-T^2A^) were generated by PCR based mutagenesis. These 3xFLAG-CENP-T mutants were then cloned into the plasmid containing approximately 2 kb homology arm around the start codon of *CENP-T* with the neomycin resistance gene cassette and hisD gene cassette by In-Fusion^®^ HD (TAKARA) (pBS_CENP-T N-KI_3xFLAG-CENP-T^WT^, _3xFLAG-CENP-T^Δ161–216^,_3xFLAG-CENP-T^RR-AA^, _3xFLAG-CENP-T^IFV-RRR^, or _3xFLAG-CENP-T^2A^_NeoR or _HisD).

### Generation of cell lines

To express the GFP-Dsn1 constructs under control of the endogenous *Dsn1* promoter, pBS_Dsn1-N KI_GFP-Dsn1^WT^, _GFP-Dsn1^4A^, or _GFP-Dsn1^Δ93–114^_PuroR and pBS_Dsn1-N KI_GFP-Dsn1^WT^, _GFP-Dsn1^4A^, or _GFP-Dsn1^Δ93–114^_EcoGPT were transfected with pX330_ggDsn1 ^23^ using Neon Transfection System (Thermo Fisher) with 6 times of pulse (1400 V, 5 msec) in CL18 (wild-type) cells, CENP-C^Δ73^cells ^23^, and CENP-C^Δ73^ cells expressing mScarlet-mAID-CENP-T.

To express OsTIR1-T2A-BSR_IRES_mScarlet-mAID-CENP-T construct under control of the CMV promoter from the endogenous PGK locus, pBS_PGK N-KI_pCM-OsTIR1-T2A-BSR-IRES-mScarlet-mAID-CENP-T was transfected with pX335_ggPGK ^35^ using Neon Transfection System (Thermo Fisher) with 6 times of pulse (1400 V, 5 msec) in CENP-C^Δ73^cells ^23^.

To express 3xFLAG-CENP-T constructs under control of the endogenous *CENP-T* promoter, pBS_CENP-T N-KI_ 3xFLAG-CENP-T^WT^, 3xFLAG-CENP-T^Δ161–216^, 3xFLAG-CENP-T^RR-AA^, 3xFLAG-CENP-T^IFV-RRR^, or 3xFLAG-CENP-T^2A^ _NeoR and pBS_CENP-T N-KI_ 3xFLAG-CENP-T^WT^,3xFLAG-CENP-T^Δ161–216^, 3xFLAG-CENP-T^RR-AA^, 3xFLAG-CENP-T^IFV-RRR^, or 3xFLAG-CENP-T^2A^ _HisD were transfected with pX335_ggCENP-T ^18^ using Neon Transfection System (Thermo Fisher) with 6 times of pulse (1400 V, 5 msec) in CENP-C^Δ73^/GFP-Dsn1^WT^ cells expressing mScarlet-mAID-CENP-T, CENP-C^Δ73^/GFP-Dsn1^4A^ cells expressing mScarlet-mAID-CENP-T.

The transfected cells were selected in the medium containing appropriate selection drugs (0.5 µg/mL puromycin (TAKARA) for the selection of PuroR expression, 25 µg/mL mycophenolic acid (Wako) and 125 µg/mL xanthine (SIGMA) for the selection of EcoGPT expression, 25 µg/mL Blasticidin S hydrochloride (Wako) for the selection of BSR expression, and L-histidinol for the selection of HisD expression) in 96 well plates to isolate single colonies.

### Structure prediction

Structures were predicted with AlphaFold2-Multimer through ColabFold using MMseqs2 and default settings ^36^. The structural figures were prepared using UCSF ChimeraX.

### Immunoblot

DT40 cells were collected, washed with cold PBS, and suspended in 1xLaemmli Sample Buffer (LSB; 62.5 mM Tris-HCl (pH 6.8), 10 % Glycerol, 2% SDS, 5% 2-mercaptoethanol, bromophenol blue) (final concentration 1×10^4^ cells/µL). Following the sonication, the lysate was heated at 96°C for 5 min.

The collected samples (1×10^5^ cells) were separated by SDS-PAGE (SuperSep Ace, 5–20% (Wako) and transferred onto PVDF membranes (Immobilon®-P (Merck)). The membrane was probed with primary antibody diluted with Signal Enhancer Hikari (Nacalai Tesque) at 4°C for overnight. After washing with 0.1% TBST (TBS, 0.1% Tween 20) for 15 min, the membrane was probed with secondary antibody diluted with Signal Enhancer Hikari (Nacalai Tesque) at room temperature for 1 h. After washing with 0.1% TBST for 15 min, the signals were detected using ECL Prime (GE Healthcare) and visualized by ChemiDoc Touch imaging system (Bio-Rad). The Image processing was performed using Image Lab 5.2.1 (Bio-Rad) and Photoshop CC (Adobe).

Primary antibodies used in immunoblot analyses were rabbit anti-chicken CENP-T ^5^, rabbit anti-chicken CENP-C ^37^, rabbit anti-chicken Dsn1 ^18^, mouse anti-FLAG M2 (Sigma), rat anti-RFP (Chromotek), and mouse anti-α-tubulin (Sigma). Secondary antibodies used in immunoblot analysis were horseradish peroxidase-conjugated (HRP)-conjugated goat anti-rabbit IgG and HRP-conjugated rabbit anti-mouse IgG (Jackson ImmunoResearch).

### Immunoprecipitation

For 3xFLAG-CENP-T immunoprecipitation, cells expressing mScarlet-mAID-CENP-T and 3xFLAG-CENP-T were cultured with 500 µM IAA for 12 h and with 100 ng/mL nocodazole for the last 10h. These cells were collected, washed with cold PBS twice, suspended in Lysis buffer (20 mM Hepes-NaOH (pH 7.4), 150 mM NaCl, 0.1% NP40, 5 mM 2-mercaptoethanol, 1xcomplete EDTA-free proteinase inhibitor (Roche), and 1xPhosSTOP (SIGMA) (final: 2×10^8^ cells/mL) and sonicated. The lysate was treated with TURBO nuclease (50 unit/mL) for 30 min, clarified by centrifugation, and the supernatant was incubated with Protein-G Dynabeads (Thermo Fisher Scientific) conjugated to anti FLAG-M2 antibody at 4°C for 1 h. Proteins precipitated with anti FLAG-M2 antibody bound beads were washed with Lysis buffer three times, eluted with 2×LSB. The eluted samples were heated at 96°C for 5 min and subjected to immunoblot analysis.

### Immunofluorescence analysis

The cells were cytospan onto slide glasses or coverslips by the Cytospin3 centrifuge (Shandon). For the CENP-T staining, cells were fixed with 3% paraformaldehyde (PFA) in 250 mM HEPES-NaOH (pH 7.4) at RT for 15 min and permeabilized in 0.5% NP-40 in PBS at RT for 10 min. For the CENP-A staining, cells were fixed with 3% paraformaldehyde (PFA) in 250 mM HEPES-NaOH (pH 7.4) at RT for 30 sec and fixed with methanol at-30°C for 20 min. After blocking with 0.5% BSA in PBS for 5 min, the cells were incubated with primary antibodies diluted in 0.5% BSA in PBS at 37°C for 1 h. The cells were washed three times with 0.5% BSA in PBS, incubated with secondary antibodies diluted in 0.5% BSA in PBS at 37°C for 1 h, and washed three times with 0.5% BSA in PBS. The cells were post-fixed for 10 min. DNA was stained with 100 ng/mL DAPI in PBS for 20 min. The stained cells were mounted with VECTASHIELD Mounting Medium (Vector Laboratories).

Primary antibodies used in immunofluorescence analysis were rabbit anti-chicken CENP-T ^5^, mouse anti-FLAG-M2 (Sigma), mouse anti-α-tubulin (Sigma). Secondary antibodies used in immunofluorescence analysis were FITC-conjugated goat anti-rabbit IgG F(ab’)2, FITC-conjugated goat anti-mouse IgG, FITC-conjugated goat anti-rat IgG, Cy3-conjugated mouse anti-rabbit IgG, Cy3-conjugated goat anti-mouse IgG, and Alexa647-conjugated goat anti-mouse IgG (Jackson ImmunoResearch).

### Image acquisition

Immunofluorescence images were acquired every 0.2 μm intervals of z-slice using a Zyla 4.2 sCMOS camera (ANDOR) mounted on a Nikon Eclipse Ti inverted microscope with an objective lens (Nikon; Plan Apo lambda 100x/1.45 NA) with a spinning disk confocal scanner unit (CSU-W1; Yokogawa) controlled with NIS-elements (Nikon). The images in the figures are the maximum intensity projection or sum intensity projection (chromosome spread samples) of the Z-stack generated with Fiji ^38^.

### Quantification and statistical analysis

The fluorescence signal intensities of GFP-Dsn1 on kinetochores and background signals in non-kinetochore region for each sample were quantified using Imaris (Bitplane). The mean of fluorescence signal intensities on kinetochores in each cell were subtracted with mean of background signals in non-kinetochore region.

Statistical analyses were performed using GraphPad Prism (GraphPad Software).

### Chromosome alignment assay

The cells were treated with IAA (500 μM) for 12 h and incubated with MG132 (10 μM) for the last 2 h and harvested. The harvested cells were fixed and immunostained with anti-CENP-A antibody as above and mounted with VECTASHIELD Mounting Medium. The cells images were acquired by the confocal microscopy with spinning-disc. The chromosome alignment was analyzed according to a previous report ^39^. Briefly, The XY positions of CENP-A signals in mitotic cells were acquired using Imaris software and plotted on a two-dimensional plane. Scatter plot analysis was performed with confidence ellipses using Real Statistics in Excel (www.real-statistics.com). Chromosome alignment was assessed by calculating the ratio of the semi-major axis to the semi-minor axis of the ellipse and subtracting 1 to obtain the alignment value.

### Flow cytometry analysis

The cells were treated with IAA (500 μM) for 12 h and incubated with 20 μM BrdU (5-bromo-2-deoxyuridine) for 20 min and harvested. The harvested cells were washed with the ice-cold PBS, fixed with 70% ethanol at –30 °C. The fixed cells were washed with 1% BSA/PBS, incubated in 4 N HCl with 0.5% Triton X-100 at RT for 30 min, and washed with 1% BSA/PBS three times. The cells were incubated with an anti-BrdU antibody (BD, 347580) at RT for 1 h and washed with 1% BSA/PBS two times. Then, the cells were incubated with FITC-conjugated anti-mouse IgG (Jackson ImmunoResearch, 115-095-003; at 1:20 dilution in 1% BSA/PBS) at RT for 30 min, washed with 1% BSA/PBS and stained DNA with propidium iodide (10 μg/mL) in 1% BSA/PBS at 4 °C overnight. The stained cells were applied to a Flow cytometry (BD, FACS Canto II). The cell cycle gates were manually adjusted and the percentages of each cell cycle stage (subG1, G1, S, G2/M phase or other cell) were calculated.

### LC-MS/MS analysis

For 3xFLAG-CENP-T immunoprecipitation, cells expressing 3xFLAG fused CENP-T, spot-fused Dsn1, and CENP-C^Δ73^ were cultured with 100 ng/mL nocodazole for 12 h. These cells were collected, washed with cold PBS twice, and suspended in 300 μL Lysis buffer (20 mM Hepes-NaOH (pH 7.4), 150 mM NaCl, 0.1% NP40, 5 mM 2-mercaptoethanol, 1xcomplete EDTA-free proteinase inhibitor (Roche), and 1xPhosSTOP (SIGMA) (final: 2×10^8^ cells/mL)). The suspension was then sonicated. The lysate was treated with TURBO nuclease (50 units/mL) for 30 min, clarified by centrifugation, and the supernatant was incubated with Protein-G Dynabeads (Thermo Fisher Scientific) conjugated to anti FLAG-M2 antibody at 4°C for 2 h. Proteins precipitated with anti FLAG-M2 antibody-bound beads were washed with 30 μL Lysis buffer five times and eluted with 25 μL Elution buffer (150 μg/mL 3xFLAG-peptide in Lysis buffer) four times.

For LC-MS/MS analysis, eluted samples underwent reduction with 10 mM Tris(2-carboxyethyl)phosphine Hydrochloride (TCEP) and alkylation via incubation with 55 mM iodoacetamide (IAA) for 30 min at room temperature in the dark. Trypsin digestion and sample cleanup were performed using the SP3 method ^40^.

Mass spectra were acquired with an Orbitrap Eclipse (Thermo Fisher Scientific) coupled to a nanoflow UHPLC system (Vanquish; Thermo Fisher Scientific). The peptide mixtures were loaded onto a C18 trap column (PepMap Neo Trap Cartridge, ID 0.3 mm x 5 mm, particle size 5 μm, Thermo Fisher Scientific) and fractionated through the C18 analytical column (Aurora, ID 0.075 × 250 mm, particle size 1.7 μm, IonOpticks). The peptides were eluted at a flow rate of 300 nl/min using the following gradient: 0% to 2% solvent B over 1 minute, 2% to 5% over 2 minutes, 5% to 16% over 19.5 minutes, 16% to 25% over 10 minutes, 25% to 35% over 4.5 minutes, a sharp increase to 95% over 4 minutes, hold at 95% for 5 minutes, and finally re-equilibration at 5%. Solvent A and B compositions were 100% H2O, 0.1% Formic acid and 100% Acetonitrile, 0.1% Formic acid, respectively. The Orbitrap operated in a data-dependent mode with a 3-second cycle time. The MS1 scan was collected at 60,000 resolution, the mass range 375-1500 m/z, using a standard AGC and maximum injection time of 50 ms. MS/MS was triggered from precursors with intensity above 20,000 and charge states 2-7. Quadrupole isolation width was 1.6 m/z, with normalized HCD energy of 30%, and resulting fragment ions recorded in Orbitrap. MS1 scan was collected at 15,000 resolution with standard AGC target and maximum injection time of 22 ms. Dynamic exclusion was set to 20 seconds.

The raw data files were searched against the Gallus gallus dataset (Uniprot Proteome UP000000539, downloaded on 20230703) with the common Repository of Adventitious Proteins (cRAP, ftp://ftp.thegpm.org/fasta/cRAP) and 3xFlag sequence for contaminant protein identification, using Proteome Discoverer 2.5 software (Thermo Fisher Scientific) with MASCOT ver.2.8 search engine, with a false discovery rate (FDR) set at 0.01. The number of missed cleavages sites was set as 2. Carbamidomethylation of cysteine was set as a fixed modification. Oxidation of methionine, phosphorylation of serine, threonine and tyrosine, and acetylation of protein N-termini were set as variable modifications.

## Data availability

All data supporting the findings of this study are available within the paper and supplementary information or from the corresponding author upon request.

---

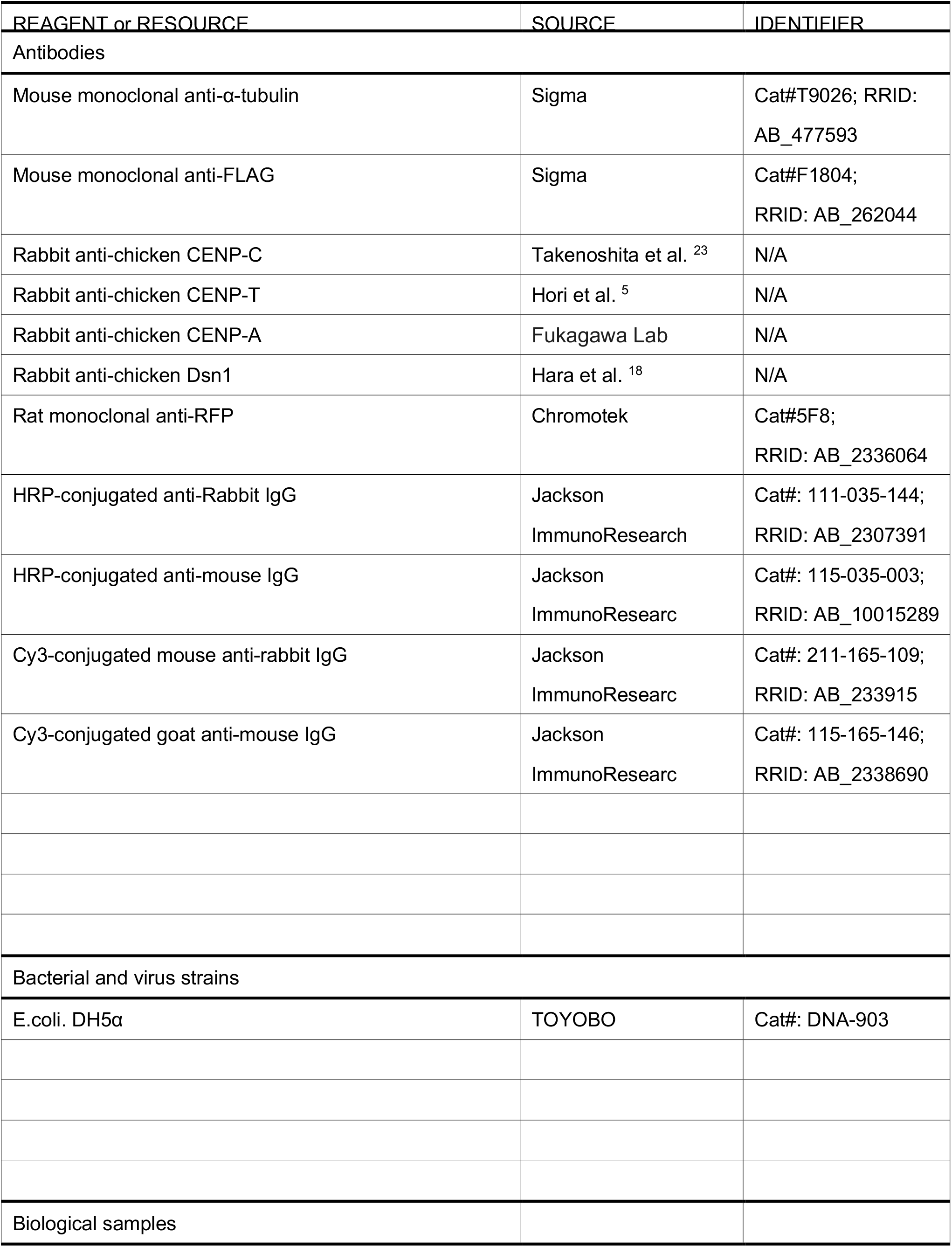
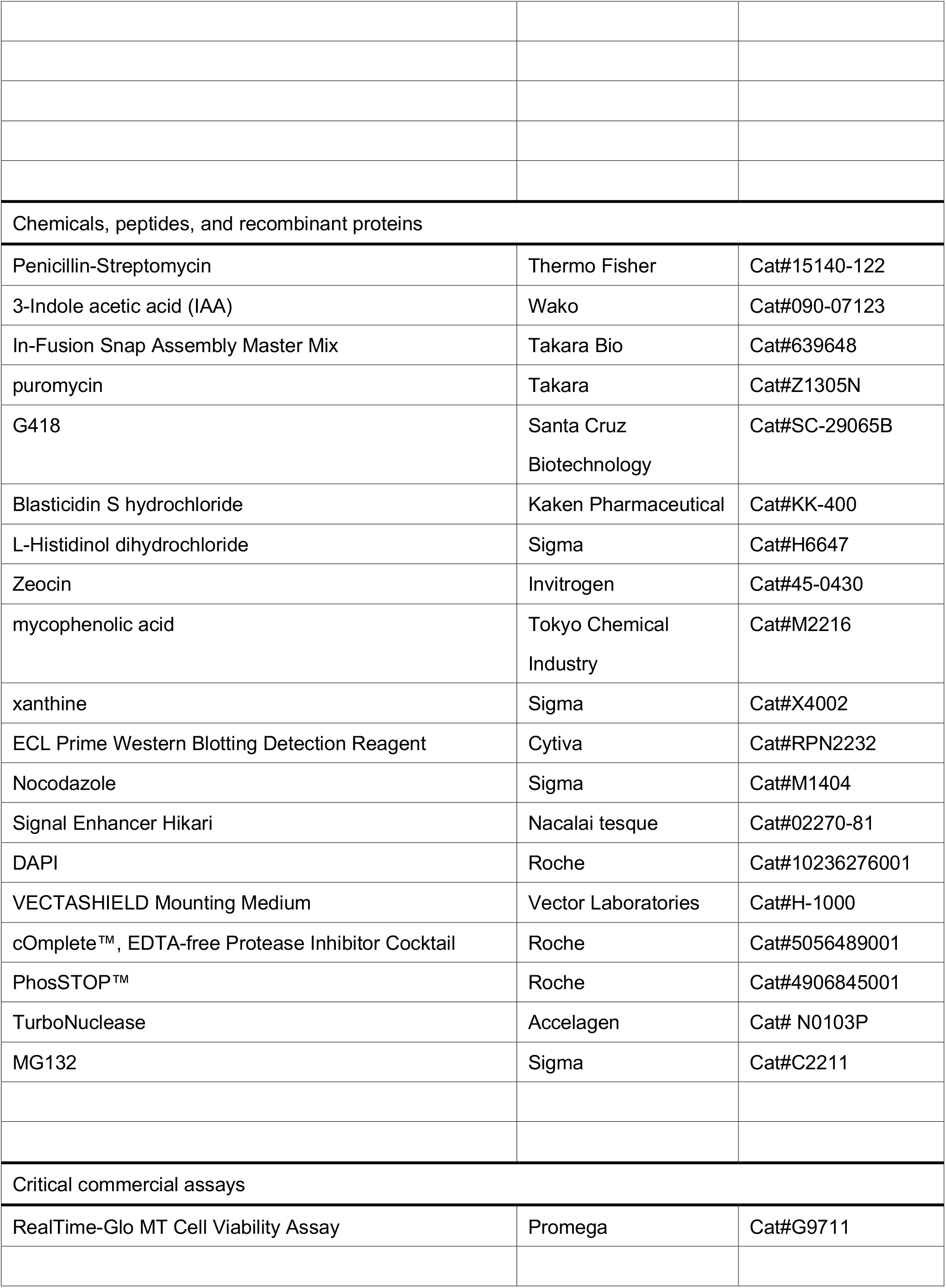
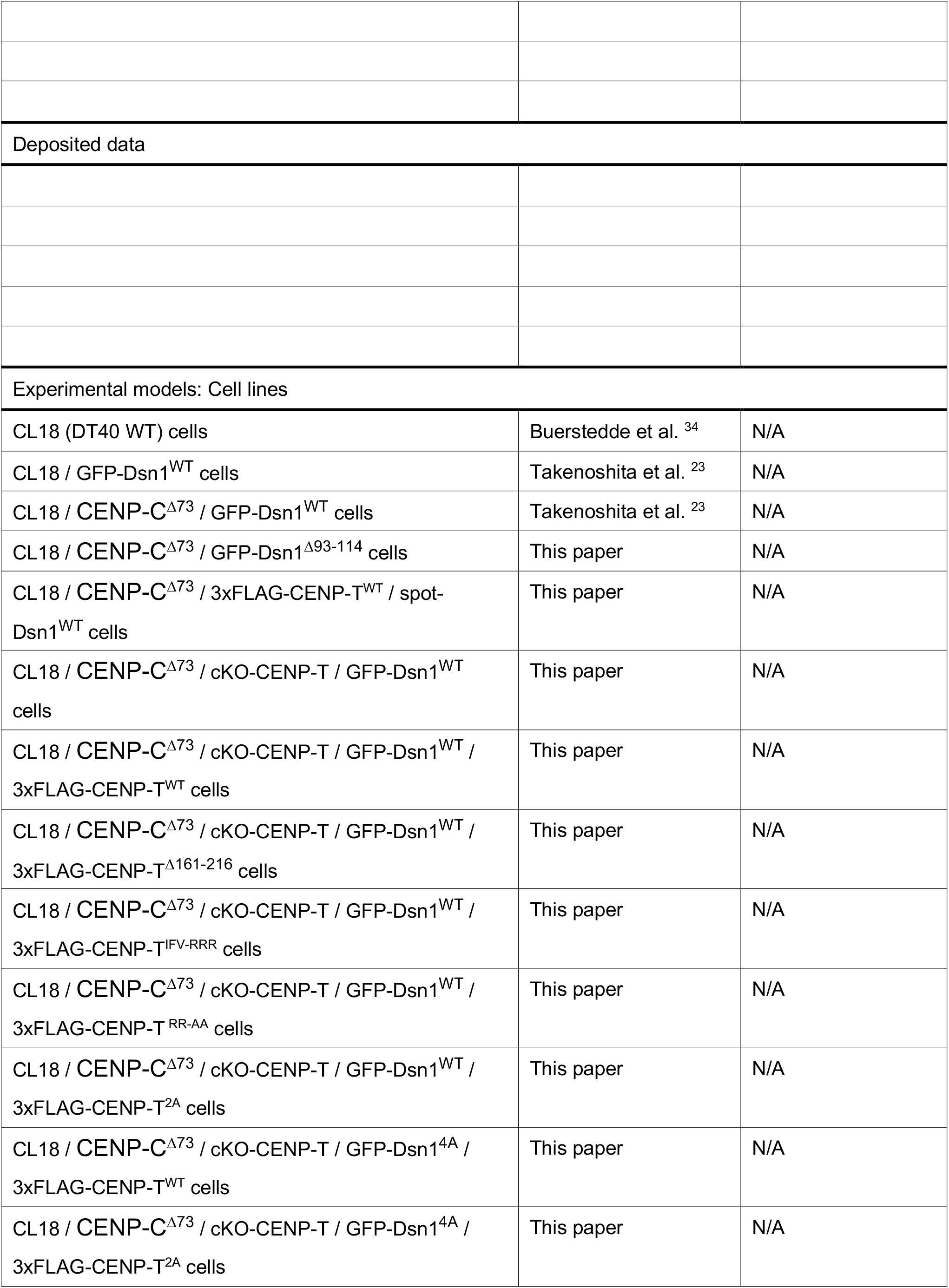
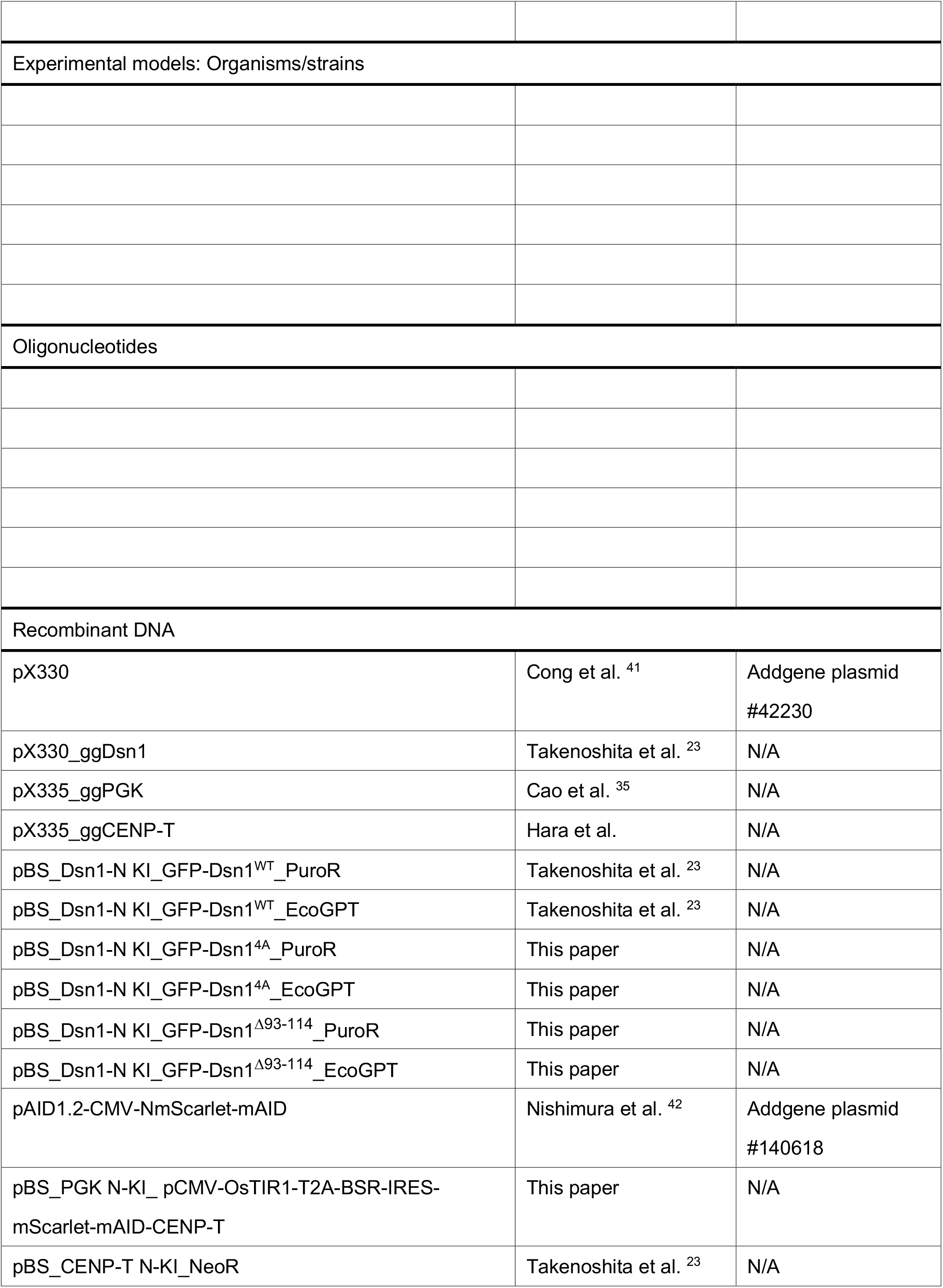
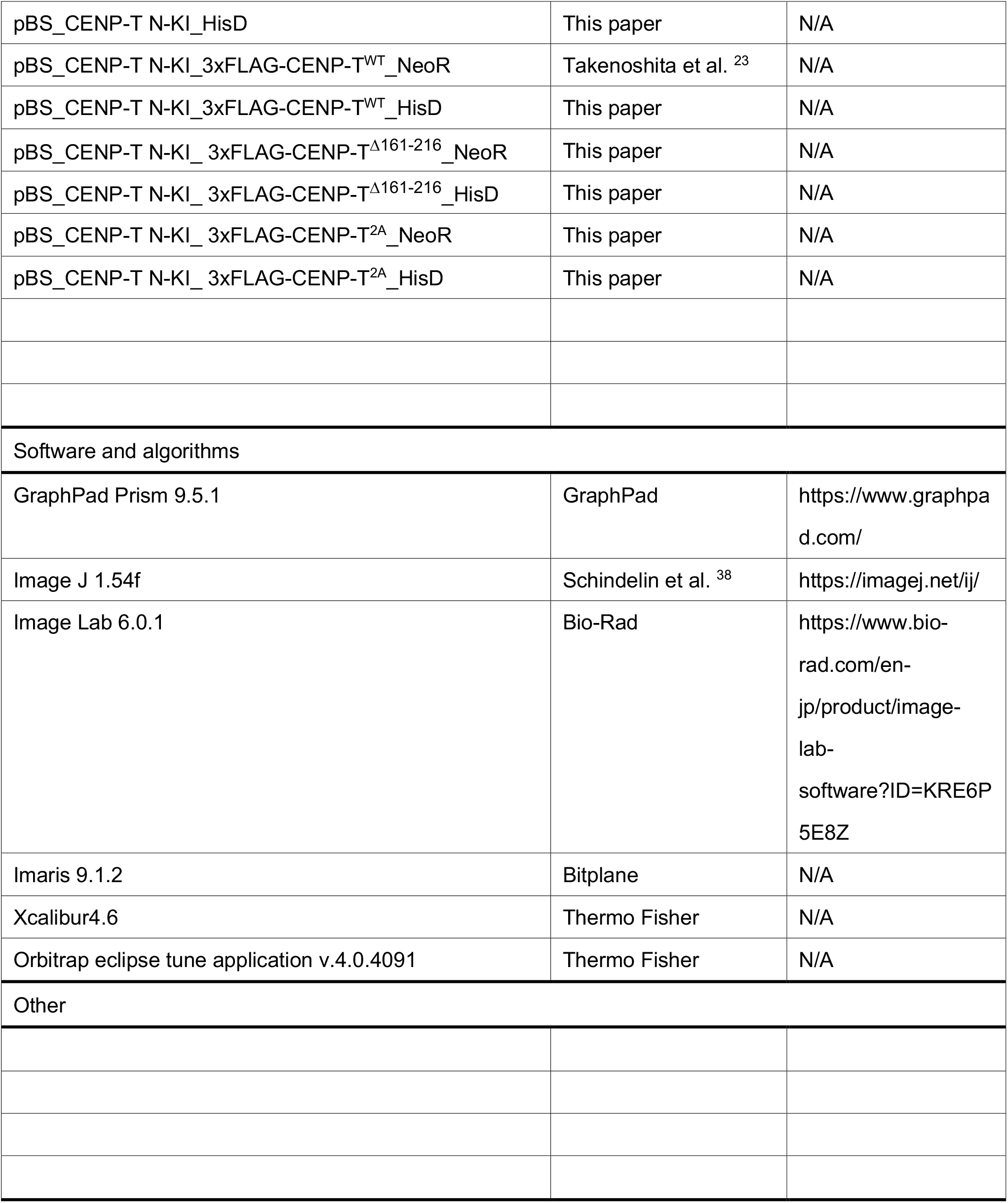
Key resources table.

## Acknowledgments

The authors are very grateful to Y. Kubota and R. Fukuoka for their technical assistance. This work was supported by CREST of JST (21460153), JSPS KAKENHI Grant Numbers 20H05389, 21H05752, 22H00408, 22H04692, and 24H02281 to TF, JSPS KAKENHI Grant Numbers 22K20630 and 23K14180 to YT, JSPS KAKENHI Grant Numbers 21H02461, 22H04672, and 24K02005, Takeda Science Foundation and Daiichi Sankyo Foundation of Life Science to MH, JSPS KAKENHI Grant Number 24K09340 to MA.

## Author contributions

Conceptualization: YT, TF; Investigation: YT, MH, RN, MA; Formal analysis: YT, RN; Resources: MH; Data curation: YT; Writing-original draft: TF; Writing-review & editing: TF, YT; Project administration: TF; Funding acquisition: TF, YT, MH, MA.

## Competing Interests

The authors declare no competing interests.

Correspondence and requests for materials should be addressed to TF.

## Supplementary Figure legends

**Figure S1.**
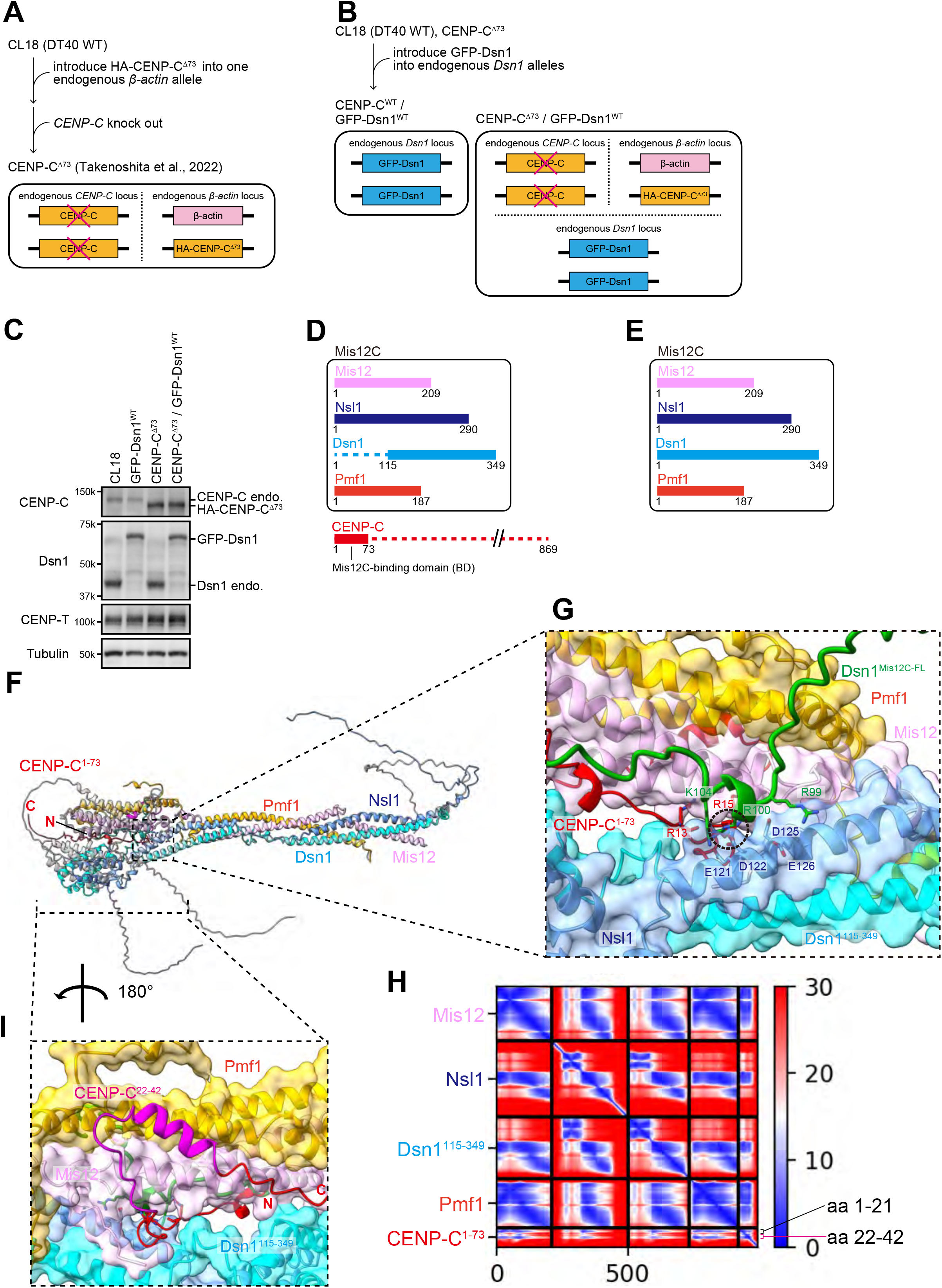
Experimental strategy of generation of CENP-C^Δ73^ cells expressing GFP-Dsn1. (A) Schematic representation of the strategy for generating CENP-C knockout DT40 cells expressing HA-CENP-C^Δ73^ (CENP-C^Δ73^ cells). The endogenous CENP-C gene was knocked out after introducing HA-CENP-C^Δ73^ into the endogenous β-actin locus via genome editing. (B) Schematic representation of the strategy for generating CENP-C^Δ73^ cells expressing GFP-Dsn1. GFP-Dsn1 was introduced into the endogenous Dsn1 locus in CENP-C^Δ73^ cells (CENP-C^Δ73^/GFP-Dsn1 cells). GFP-Dsn1 was also introduced into the endogenous Dsn1 locus in CL18 (wild-type DT40) cells. (C) Immunoblot analysis of CL18 or CENP-C^Δ73^ cells expressing GFP-Dsn1 with the indicated antibodies. The endogenous CENP-C and HA-CENP-C^Δ73^ proteins were detected using an anti-CENP-C antibody. The endogenous Dsn1 and GFP-Dsn1 proteins were detected using an anti-Dsn1 antibody. The endogenous CENP-T proteins were detected using an anti-CENP-T antibody. Alpha-tubulin (Tubulin) served as the loading control. (D) Schematic representation of chicken CENP-C and chicken Mis12C used for structural prediction of the CENP-C-Mis12C complex. The full length of Mis12 (209 aa), Nsl1 (209 aa), Pmf1 (187 aa), the region of Dsn1 aa 116–349, and the region of CENP-C aa 1–73 were used. (E) Schematic representation of the chicken Mis12C used for structural prediction of Mis12C. Mis12 (209 aa), Nsl1 (290 aa), Dsn1 (349 aa), and Pmf1 (187 aa). (F) The CENP-C-Mis12C complex predicted structure (D) overlaid on the Mis12C structure (E), both displayed as cartoons. The CENP-C-Mis12C complex predicted structure is color-coded, as indicated in the figure. The Mis12C predicted structure is colored gray, while the region of Dsn1 aa 94–114 is colored green. (G) The transparent surface model of the CENP-C-Mis12C predicted structure is overlaid on a cartoon model, while the CENP-C is displayed only as a cartoon. The Dsn1 aa 94–114 in the Mis12C is displayed as a cartoon. The key residues of the interactions between Mis12C and CENP-C or Mis12C and the Dsn1 aa 91–114 region are labeled. Models are color-coded as in (F). The steric clash region is highlighted with a black dashed circle. (H) The predicted Aligned Error (PAE) plot of the rank_1 model of the CENP-C-Mis12C complex shown in (G). CENP-C appears to bind with Mis12C via two binding regions: aa 1–21 and aa 22–42. The CENP-C aa 1–21 contain R13 and R15, which bind to the acidic amino acids of Nsl1, as shown in (G). CENP-C aa 22–42 appears to bind with Mis12 and Pmf1, as shown in (I). (I) Overview of the CENP-C-Mis12C interaction via the CENP-C aa 22–42. Models are displayed as in (G). The CENP-C aa 22–42 are colored in magenta.

**Figure S2.**
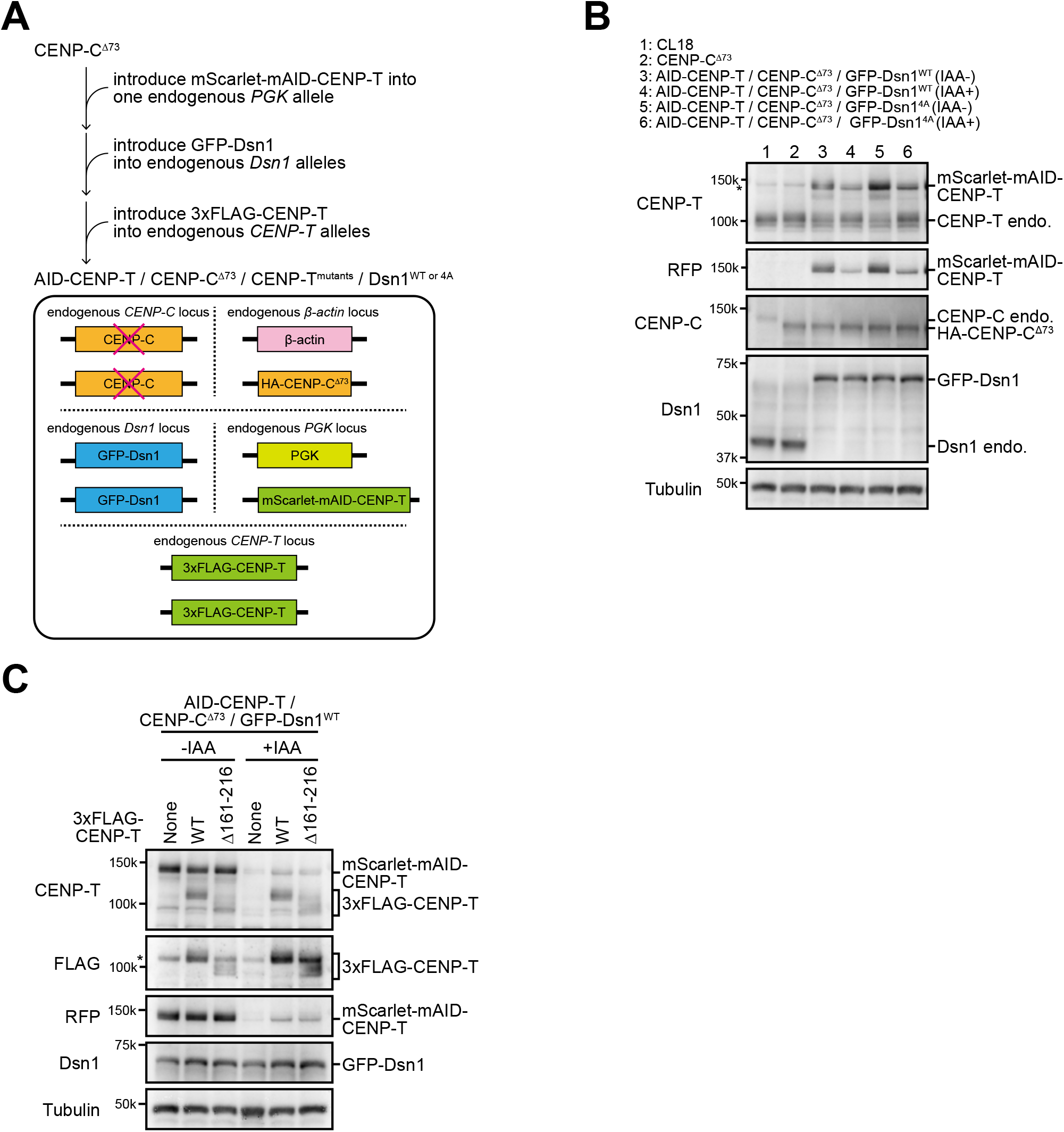
Experimental strategy for generating several variations of AID-based CENP-T conditional knockdown DT40 cells. (A) Schematic representation of a strategy for generating AID-based CENP-T conditional knockdown DT40 cells expressing HA-CENP-C^Δ73^, GFP-Dsn1, and 3xFLAG-CENP-T. First, the mScarlet-mAID-CENP-T construct containing OsTIR1 was introduced into the endogenous *PGK* locus in CENP-C^Δ73^ cells. GFP-Dsn1^WT^ or GFP-Dsn1^4A^ was introduced into the endogenous *Dsn1* locus, and 3xFLAG-fused various CENP-T (wild-type or various mutants) was introduced into the endogenous *CENP-T* locus. (B) Immunoblot analysis of CL18 (DT40 WT) cells or CENP-C^Δ73^ cells expressing GFP-Dsn1^WT^ or GFP-Dsn1^4A^ and mScarlet-mAID-CENP-T with indicated antibodies. Cells were treated with IAA for 12 h (lanes 4 and 6). The endogenous CENP-T and mScarlet-mAID-CENP-T proteins were detected using an anti-CENP-T antibody. mScarlet-mAID-CENP-T proteins were also detected using an anti-RFP antibody. Endogenous CENP-C and HA-CENP-C^Δ73^ proteins were detected using an anti-CENP-C antibody. Endogenous Dsn1 and GFP-Dsn1 proteins were detected using an anti-Dsn1 antibody. Alpha-tubulin (Tubulin) was used as the loading control. Asterisk indicates non-specific bands. (C) Immunoblot analysis of AID-based CENP-T conditional knockdown DT40 cells expressing HA-CENP-C^Δ73^, GFP-Dsn1, and 3xFLAG-CENP-T mutants with the indicated antibodies. Cells were cultured in either the absence or presence of IAA for 12 h. The 3xFLAG-CENP-T mutants and mScarlet-mAID-CENP-T proteins were detected using an anti-CENP-T antibody. The 3xFLAG-CENP-T mutants were detected using an anti-FLAG antibody. mScarlet-mAID-CENP-T proteins were detected using an anti-RFP antibody. The GFP-Dsn1 proteins were detected using an anti-Dsn1 antibody. Alpha-tubulin (Tubulin) was used as the loading control. Asterisk indicates non-specific bands.

**Figure S3.**
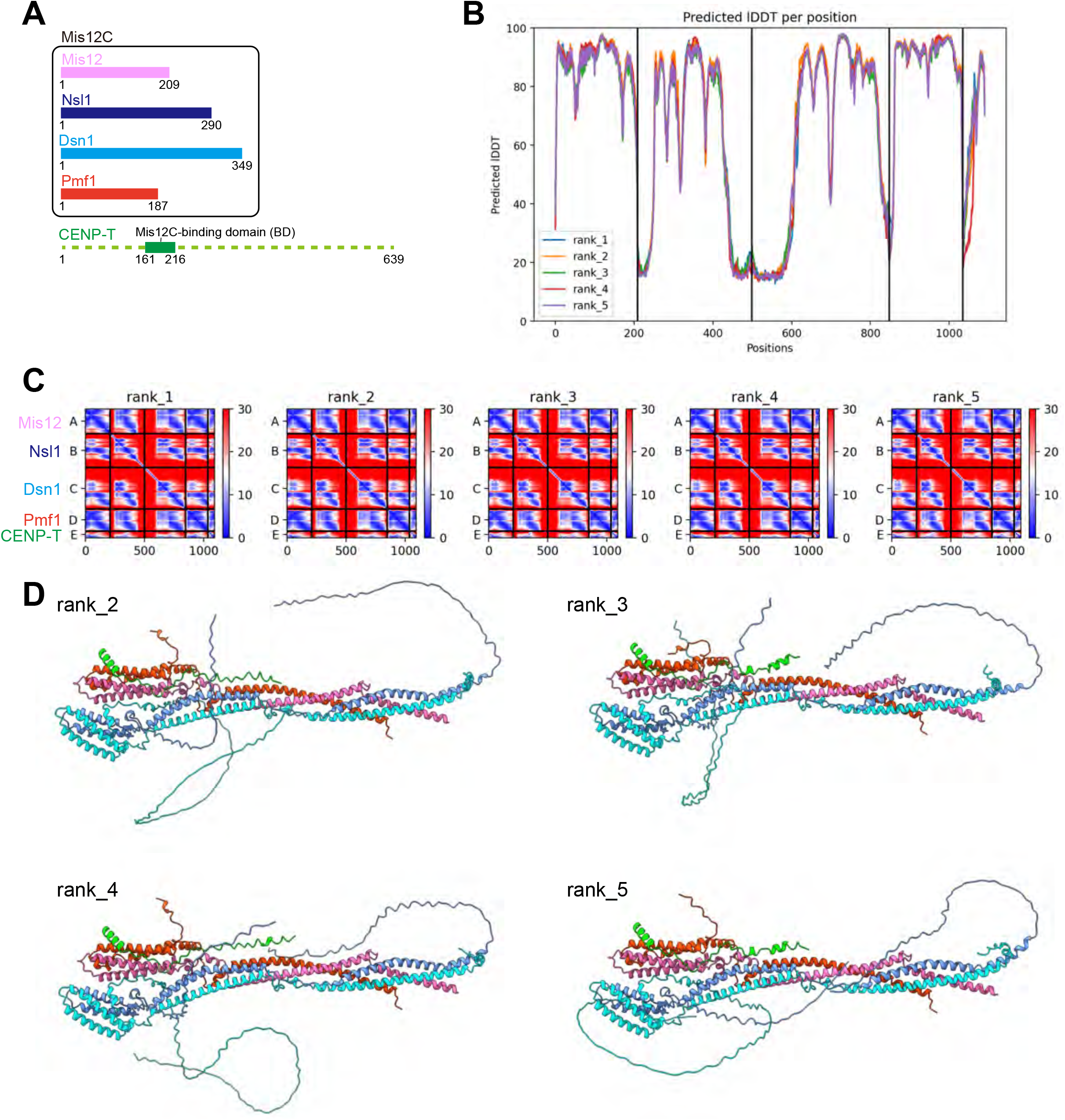
AlphaFold2 prediction of Mis12C in complex with CENP-T. (A) Schematic representation of CENP-T (639 aa) and Mis12C components: Mis12 (209 aa), Nsl1 (290 aa), Dsn1 (349 aa), and Pmf1 (187 aa). The full length of the Mis12C components and the region of CENP-T aa 161–216 were used for this prediction. (B and C) pLDDT score plot (B) and Predicted Aligned Error (PAE) plot (C) of the rank_1 model are shown in Figure 2A; rank_2, 3, 4, and 5 models are shown in (D). (D) AlphaFold2 predicted structures with ranks 2, 3, 4, and 5.

**Figure S4.**
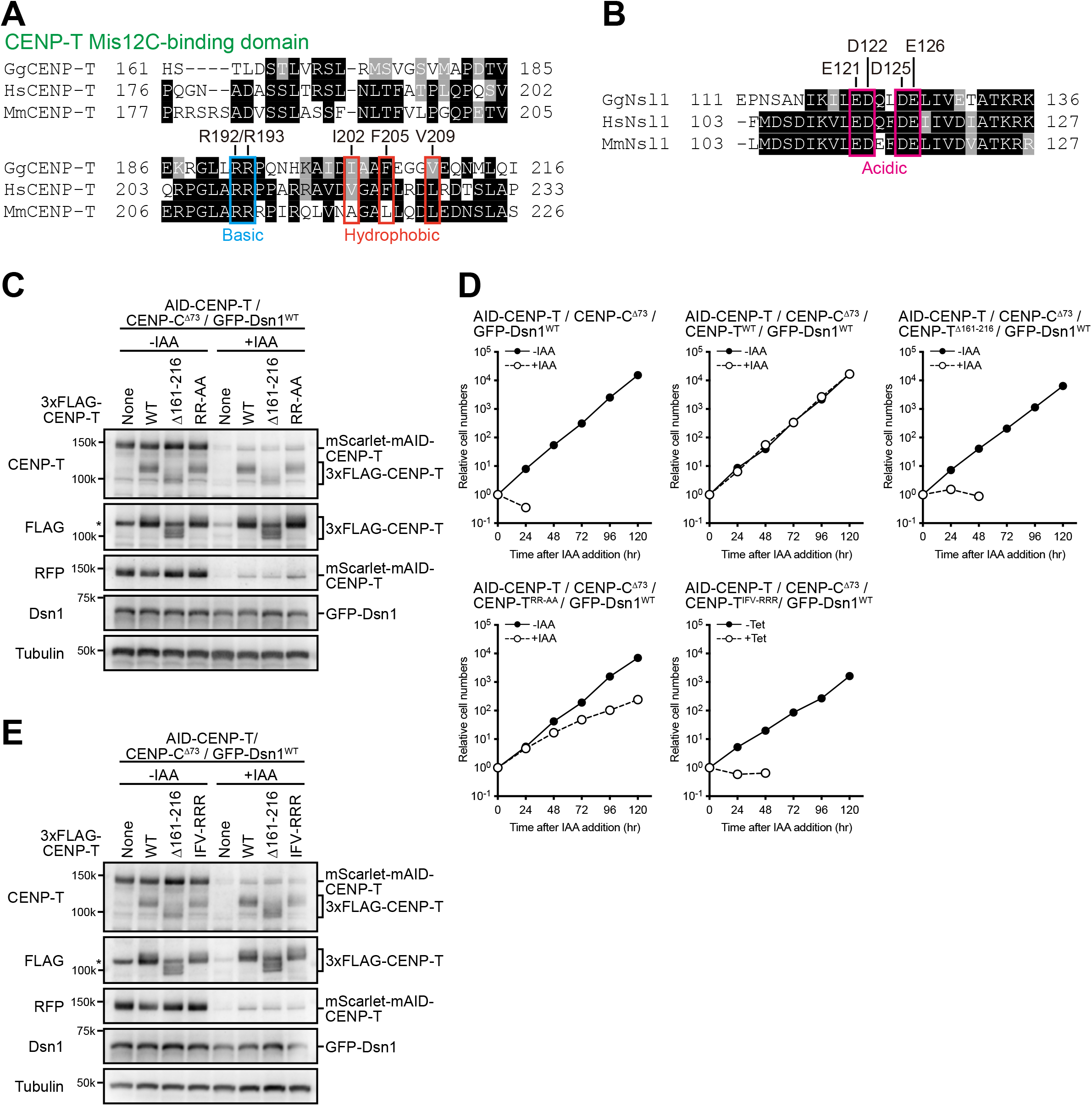
Characterization of AID-based CENP-T knockdown CENP-C^Δ73^ cells expressing various CENP-T mutants. (A) Amino acid alignment of a Mis12C-binding domain of CENP-T among various species. Conserved arginine residues (R192 and R193 in chicken CENP-T) and hydrophobic residues (I202, F205, and V209 in chicken CENP-T) are highlighted. (B) Amino acid alignment of an acidic region of Nsl1 among various species. Conserved acidic residues (E121, D122, D125, and E126 in chicken Nsl1) are highlighted. (C) Immunoblot analysis of AID-based CENP-T conditional knockdown DT40 cells expressing HA-CENP-C^Δ73^, GFP-Dsn1, and 3xFLAG-CENP-T mutants with the indicated antibodies. Cells were cultured in either the absence or presence of IAA for 12 h. The 3xFLAG-CENP-T mutants and mScarlet-mAID-CENP-T proteins were detected using an anti-CENP-T antibody. The 3xFLAG-CENP-T mutants were detected using an anti-FLAG antibody. mScarlet-mAID-CENP-T proteins were detected using an anti-RFP antibody. The GFP-Dsn1 proteins were detected using an anti-Dsn1 antibody. Alpha-tubulin (Tubulin) was used as the loading control. Asterisk indicates non-specific bands. (D) Growth analysis of AID-based CENP-T conditional knockdown DT40 cells expressing HA-CENP-C^Δ73^, GFP-Dsn1, and 3xFLAG-CENP-T mutants in the absence or presence of IAA. Cell numbers were normalized to those at 0 h for each cell line. (E) Immunoblot analysis of AID-based CENP-T conditional knockdown DT40 cells expressing HA-CENP-C^Δ73^, GFP-Dsn1, and 3xFLAG-CENP-T mutants with the indicated antibodies. Cells were cultured in either the absence or presence of IAA for 12 h. Each protein was detected with an antibody, as described in (C).

**Figure S5.**
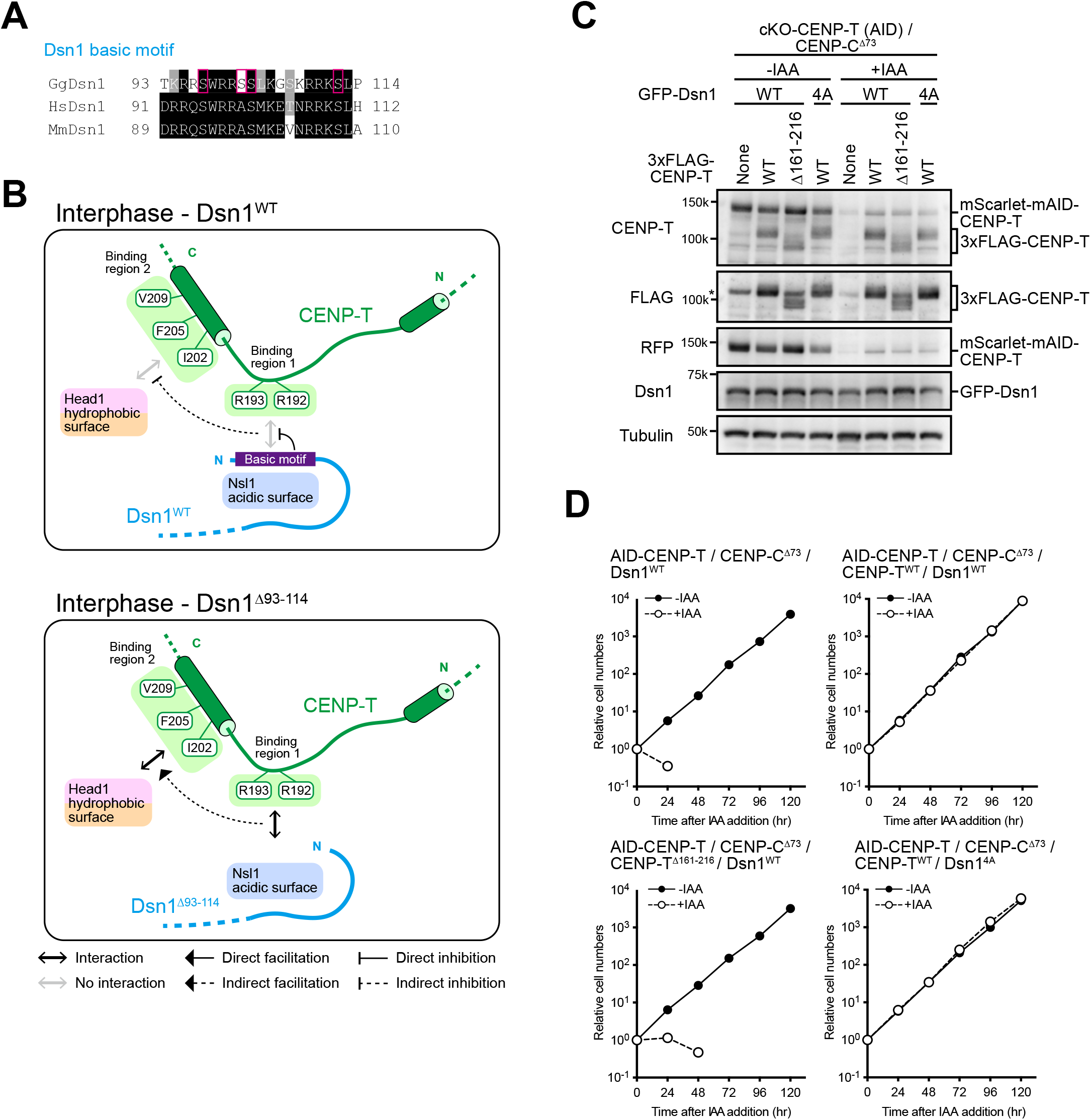
Characterization of AID-based CENP-T knockdown CENP-C^Δ73^ cells expressing Dsn1 mutants. (A) Amino acid alignment of the Dsn1 basic motif among various species. Serine residues within the Aurora B consensus motif (S97, S101, S102, and S112) were substituted to alanine in Dsn1^4A^. (B) A model of the inhibition mechanisms of the CENP-T-Mis12C interaction in interphase, based on the phenotype of CENP-C^Δ73^/Dsn1^Δ93–114^ cells. The Dsn1 basic motif masks the Nsl1 acidic surface, inhibiting interaction with the binding region 1 of CENP-T. This inhibition causes the defects of the interaction between the hydrophobic surface of head1 in Mis12C and binding region 2 of CENP-T. When the basic motif of Dsn1 is deleted, binding region 1 of CENP-T binds to Nsl1 acidic surface, leading to the interaction between hydrophobic surface of head1 in Mis12C and binding region 2 of CENP-T. (C) Immunoblot analysis of AID-based CENP-T conditional knockdown DT40 cells expressing HA-CENP-C^Δ73^, GFP-Dsn1 mutants, and 3xFLAG-CENP-T mutants with the indicated antibodies. Cells were cultured in either the absence or presence of IAA for 12 h. The 3xFLAG-CENP-T mutants and mScarlet-mAID-CENP-T proteins were detected using an anti-CENP-T antibody. The 3xFLAG-CENP-T mutants were detected using an anti-FLAG antibody. mScarlet-mAID-CENP-T proteins were detected using an anti-RFP antibody. The GFP-Dsn1^WT^ and GFP-Dsn1^4A^ proteins were detected using an anti-Dsn1 antibody. Alpha-tubulin (Tubulin) was used as the loading control. Asterisk indicates non-specific bands. (D) Growth analysis of AID-based CENP-T conditional knockdown DT40 cells expressing HA-CENP-C^Δ73^, GFP-Dsn1 mutants, and 3xFLAG-CENP-T mutants in the absence or presence of IAA. Cell numbers were normalized to those at 0 h for each cell line.

**Figure S6.**
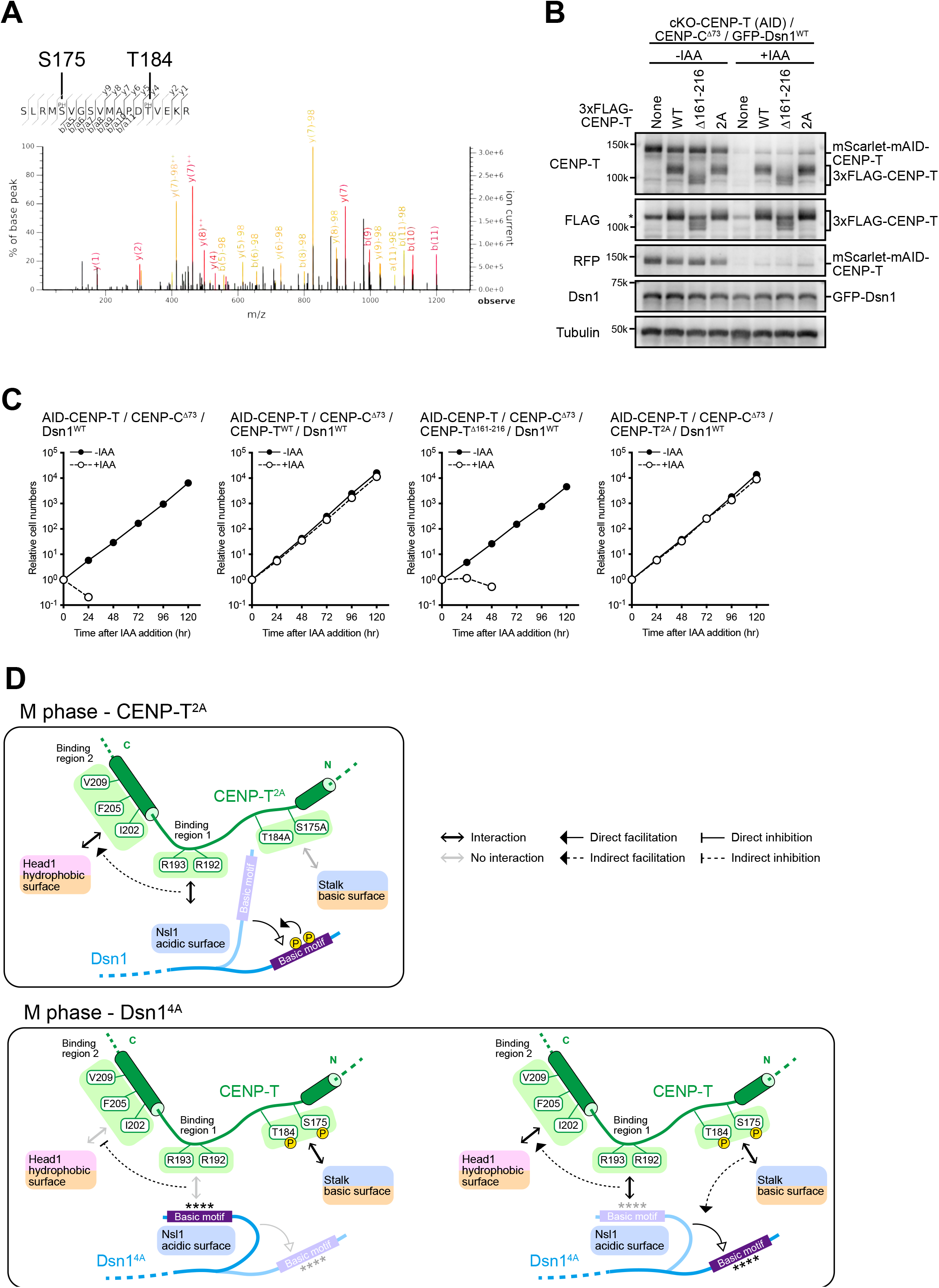
Characterization of AID-based CENP-T knockdown CENP-C^Δ73^ cells expressing CENP-T mutants. (A) Peptide spectrum of the phosphorylated CENP-T peptide. Phosphorylated amino acids, S175 and T184, are labeled in the sequence. (B) Immunoblot analysis of AID-based CENP-T conditional knockdown DT40 cells expressing HA-CENP-C^Δ73^, GFP-Dsn1, and 3xFLAG-CENP-T mutants with the indicated antibodies. Cells were cultured in either the absence or presence of IAA for 12 h. The 3xFLAG-CENP-T mutants and mScarlet-mAID-CENP-T proteins were detected using an anti-CENP-T antibody. The 3xFLAG-CENP-T mutants were detected using an anti-FLAG antibody. mScarlet-mAID-CENP-T proteins were detected using an anti-RFP antibody. The GFP-Dsn1 proteins were detected using an anti-Dsn1 antibody. Alpha-tubulin (Tubulin) was used as the loading control. Asterisk indicates non-specific band. (C) Growth analysis of AID-based CENP-T conditional knockdown DT40 cells expressing HA-CENP-C^Δ73^, GFP-Dsn1, and 3xFLAG-CENP-T mutants in the absence or presence of IAA. Cell numbers were normalized to those at 0 h for each cell line. (D) Models to explain the CENP-T-Mis12C interaction in CENP-C^Δ73^/CENP-T^2A^ cells and CENP-C^Δ73^/Dsn1^4A^ cells during M phase. In CENP-C^Δ73^/CENP-T^2A^ cells, the interaction between the phosphorylation sites of CENP-T and the basic surface in the stalk region of Mis12C is inhibited, but CENP-T still binds to Mis12C using the binding region 1 and 2. In total the binding efficiency was reduced to ∼70% levels. In CENP-C^Δ73^/Dsn1^4A^ cells, the Dsn1 basic motif masks the Nsl1 acidic surface and inhibit the interaction between binding region 1 of CENP-T and the Nsl1 acidic surface, but some population of Mis12C preclude the Dsn1 mask because some Mis12C still bind to CENP-T (Levels were reduced to ∼60%). The CENP-T-Mis12C binding phenotype of in CENP-C^Δ73^/CENP-T^2A^ cells and CENP-C^Δ73^/Dsn1^4A^ cells suggest that a coordination of CENP-T and Dsn1 phosphorylation might be necessary to completely preclude the Dsn1 mask.

**Figure S7.**
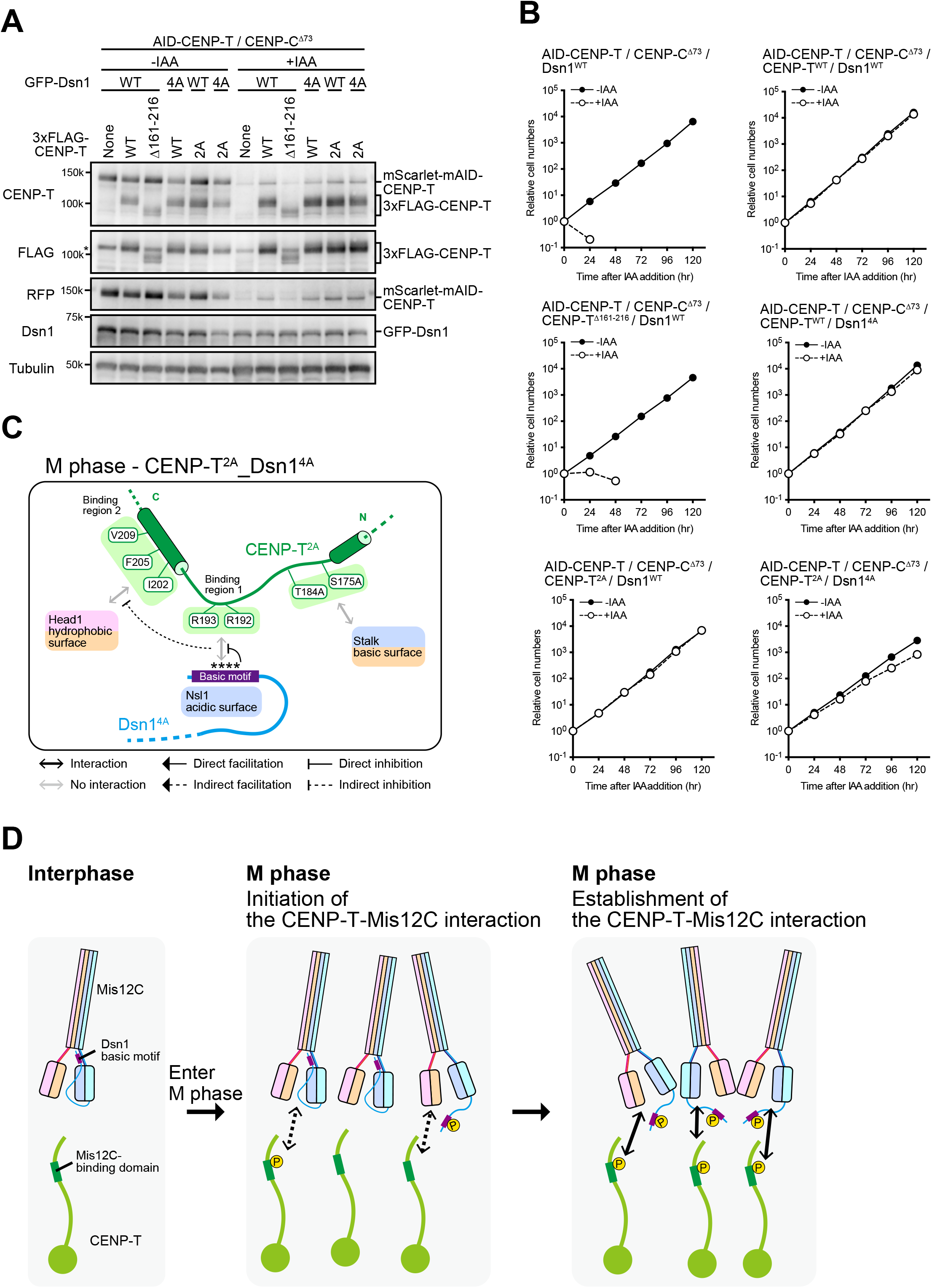
Characterization of AID-based CENP-T knockdown CENP-C^Δ73^ cells expressing CENP-T and Dsn1 mutants. (A) Immunoblot analysis of AID-based CENP-T conditional knockdown DT40 cells expressing HA-CENP-C^Δ73^, GFP-Dsn1 mutants, and 3xFLAG-CENP-T mutants with the indicated antibodies. Cells were cultured in either the absence or presence of IAA for 12 h. The 3xFLAG-CENP-T mutants and mScarlet-mAID-CENP-T proteins were detected using an anti-CENP-T antibody. The 3xFLAG-CENP-T mutants were detected using an anti-FLAG antibody. mScarlet-mAID-CENP-T proteins were detected using an anti-RFP antibody. The GFP-Dsn1^WT^ and GFP-Dsn1^4A^ proteins were detected using an anti-Dsn1 antibody. Alpha-tubulin (Tubulin) served as the loading control. Asterisk indicates non-specific band. (B) Growth analysis of AID-based CENP-T conditional knockdown DT40 cells expressing HA-CENP-C^Δ73^, GFP-Dsn1 mutants, and 3xFLAG-CENP-T mutants in the absence or presence of IAA. Cell numbers were normalized to those at 0 h for each cell line. (C) A model to explain the CENP-T-Mis12C interaction in CENP-C^Δ73^/CENP-T^2A^/Dsn1^4A^ cells during M phase. In these cells, the mask of the Dsn1 basic motif is not precluded in M phase, because both phosphorylation sites of Dsn1 and CENP-T are mutated, supporting an idea that a coordination of CENP-T and Dsn1 phosphorylation might be necessary to completely preclude the Dsn1 mask. (D) A model of the initiation and establishment of the CENP-T-Mis12C interaction after entry of M phase. Each of phosphorylation of CENP-T and Dsn1 initiates the CENP-T-Mis12C interaction, independently. Subsequently, the CENP-T and Dsn1 dual phosphorylation is coordinated to completely preclude the Dsn1 mask and the robust CENP-T-Mis12C interaction is established.

